# Parl Deficiency in Mouse Causes Coenzyme Q Depletion, Complex III Defects, and Leigh-like Syndrome

**DOI:** 10.1101/368654

**Authors:** Marco Spinazzi, Enrico Radaelli, Katrien Horré, Amaia M. Arranz, Natalia Gounko, Patrizia Agostinis, Francis Impens, Vanessa Alexandra Morais, Guillermo Lopez-Lluch, Lutgarde Serneels, Placido Navas, Bart De Strooper

## Abstract

The mitochondrial intramembrane rhomboid protease Parl has been implicated in diverse functions *in vitro*, but its physiological role *in vivo* remains unclear. Here we show that *Parl* ablation in mouse causes a striking necrotizing encephalomyelopathy similar to Leigh syndrome, a mitochondrial disease characterized by disrupted energy production. Mice with conditional Parl deficiency in the nervous system, but not in muscle, develop a similar phenotype as germline *Parl* knockouts demonstrating the vital role of Parl in neurological homeostasis. Genetic modification of two major Parl substrates, Pink1 and Pgam5, do not modify this severe neurological phenotype. *Parl^-/-^* brain mitochondria are affected by defects in Complex III activity and in coenzyme Q biosynthesis. Parl is necessary for the stable expression of Ttc19, required for Complex III activity, and of Coq4, essential in coenzyme Q biosynthesis. Thus, Parl plays a previously overseen constitutive role in the maintenance of the respiratory chain in the nervous system, and its deficiency causes progressive mitochondrial dysfunction and Leigh-like syndrome.

## INTRODUCTION

Presenilin Associated Rhomboid Like protein^1^ also called PINK1/PGAM5 Associated Rhomboid Protease^2^, both abbreviated as “PARL”, represents the only known mitochondrial member of the rhomboid family. Rhomboids are an evolutionary conserved group of intramembrane cleaving proteases and pseudo proteases involved in a variety of functions^3^. Their broad biological significance is reflected in their pathological relevance for prevalent human diseases, including cancer and neurodegenerative disease^3,4^, including Parkinson’s disease^5^.

The crucial role of PARL in cellular homeostasis is illustrated by the lethal multisystem phenotype of Parl deficient mice (*Parl^-/-^*), associated with muscle atrophy and increased apoptosis in thymus and spleen6. The faster cytochrome c release and cristae remodeling *in vitro*, the increased cell death of *Parl^-/-^* MEFs treated with apoptosis-inducing agents being rescued by overexpression of intermembrane space targeted Opa1, led to the proposal that Parl plays a crucial role in cristae remodeling and cytochrome c release during apoptosis. The authors suggested that decreased Opa1 processing by Parl was causative to these apoptotic phenotypes, although they did not exclude that Opa1 processing was also executed by other proteases. Later studies identified Oma1 and Yme1l^7^ as the proteases responsible for Opa1 processing and questioned Opa1 as a Parl substrate^8^. More recently, Parl has been implicated in the processing of other substrates in cultured cells^9–14^. Two substrates, Pink1 and Pgam5, are of particular interest as they have been implicated in the neurodegenerative disorder Parkinson’s disease^15,16^. Both accumulate in *Parl^-/-^* cells^10–12, 17^, but it is unclear whether this accumulation has a pathological effect or not^2^. A recent elegant cell biology study has led to the proposal that PARL exerts pro-apoptotic effects via misprocessing of DIABLO^14^. However, this is difficult to reconcile with the evident lethality of *Parl^-/-^* mice and the proposed anti-apoptotic, protective function of Parl^6^. Overall, the available data have led to contradictory speculations with regard to the role of PARL in mitochondrial function^6,18^, morphology^6,18,19^, mitophagy^5,20,21^ and apoptosis^6,14^ and claims have not been further substantiated *in vivo*. More importantly, the cause of death of the *Parl^-/-^* mice, and therefore the physiological role of this protease has remained unresolved. To address these questions, we decided to re-investigate Parl deficient mice. In contrast with our previous report, we find now, in addition to the previous described phenotypes in peripheral tissues, that Parl deficiency in the nervous system causes a severe dramatic necrotizing encephalomyelopathy which closely resembles pathologically Leigh syndrome, a human mitochondrial disease characterized by severe brainstem and subcortical neurodegeneration and caused by impaired energy production^22^. A similar phenotype is seen both in the full *Parl^-/-^* and in *Parl^Nes-Cre^* mice with a specific deletion of Parl in the nervous system, including the atrophy of muscle, thymus and spleen seen previously in the *Parl^-/-^* mice. This striking neurodegeneration is not associated with altered apoptosis but with massive necrosis raising the question of the underlying mechanism. We show that necrosis in *Parl^-/-^* brains is preceded by progressive mitochondrial structural changes and biochemically by defects in Complex III function and in Coenzyme Q (CoQ) biosynthesis. Thus, Parl has an essential physiological role in the maintenance of mitochondrial structure and respiratory chain function, which is severely impaired when Parl is ablated, causing Leigh-like neurodegeneration. We discuss how these novel insights affect our previous interpretations of the *Parl^-/-^* phenotype *in vivo*.

## RESULTS

### Parl deficiency in the nervous system leads to Leigh syndrome

*Parl^-/-^* mice develop normally up to the age of 40 days, after which they progressively lose weight^6^. From the age of six weeks, they show rapidly progressive locomotor impairment, paresis of the lower limbs, hunchback deformity, and dyspnea. They die before the age of eight weeks from a multisystem phenotype with atrophy of skeletal muscle, thymus and spleen ^6^, without a clear explanation. Immune deficiency is unlikely the underlying cause of death since mice thymus and spleen become atrophic only few days before death when mice are already severely affected. Moreover mice are bred in specific pathogen free conditions and do not develop opportunistic infections, leaving open the question of which essential vital function is compromised by Parl deficiency. These clinical signs led us to ask whether this striking phenotype was caused by involvement of the nervous system. Thus, we performed a detailed neuropathological analysis of *Parl^-/-^* mice. Microscopic examination of the brain and spinal cord from *Parl^-/-^* mice indeed shows a previously overlooked striking subcortical vacuolar encephalomyelopathy closely resembling Leigh syndrome (Fig. 1A-H). Leigh syndrome is a lethal mitochondrial disease characterized clinically by neurological regression and pathologically by vacuolar degeneration and necrosis of the brainstem, basal ganglia and spinal cord, associated with reactive gliosis and vascular proliferation ^23^. Vacuolization of the neuropil is first detectable in *Parl^-/-^* mice at five weeks of age, initially focally circumscribed in the brainstem and in the gray matter in the spinal cord, and then progressively extending anteriorly to hypothalamus, thalamus, deep cerebellar nuclei, and the cingulate cortex. Other areas of the brain, most notably the substantia nigra, are spared. Neuronal loss is readily detectable by the disappearance of NeuN positive cells (Fig. 1B). Neurodegeneration is accompanied by extensive astrogliosis and microgliosis, indicated by Gfap (Fig. 1C) and Iba1 (Fig. 1D) immunoreactivity. Luxol-fast blue, a staining used to visualize the white matter in the brain, shows comparable reaction in WT and *Parl^-/-^* mice (Fig. 1E). Consistent with neuronal involvement, in situ hybridization shows that *Parl* mRNA expression is particularly abundant in neurons (Fig. 1F). In advanced stages, vascular proliferation becomes evident (Fig. 1G), and symmetrical necrohemorragic foci are frequently observed at seven weeks of age in the most severely affected areas of the brainstem and spinal cord (Fig. 1H). When Parl was specifically ablated in the nervous system using a Nestin-Cre driver (*Parl^Nes-Cre^*) (Fig. 1J) a similar lethal phenotype is observed as in the germline *Parl^-/-^* mice (Fig. 1I) including the Leigh type neuropathology (Fig.1K). Apart from a 4-weeks delay in lethality and the absence of testis atrophy (Fig. 1O), these mice develop the typical *Parl^-/-^* multisystem phenotype^6^ with severe atrophy of muscle, liver, spleen (Fig.1L-N), and thymus (Fig. 1P) despite normal Parl expression in these tissues (Fig. 1J) even at late symptomatic stages. Thymus and spleen become atrophic in *Parl^Nes-Cre^* as in germline *Parl^-/-^* mice only in preterminal stages of the disease when the mice are already affected by severe neurological deficits. Conversely, deletion of Parl in striated muscle in *Parl^Ckmm-Cre^* knockout mice using *Cre* expression driven by the Creatine kinase promoter did not compromise survival beyond the age of 15 months nor led to overt locomotor deficits (n=13, Fig. 1I). Altogether, these data suggest that *Parl* deficiency in the nervous system is sufficient to recapitulate the lethal multisystem phenotype of germline *Parl^-/-^* mice, except the gonad atrophy.

**Fig. 1.**
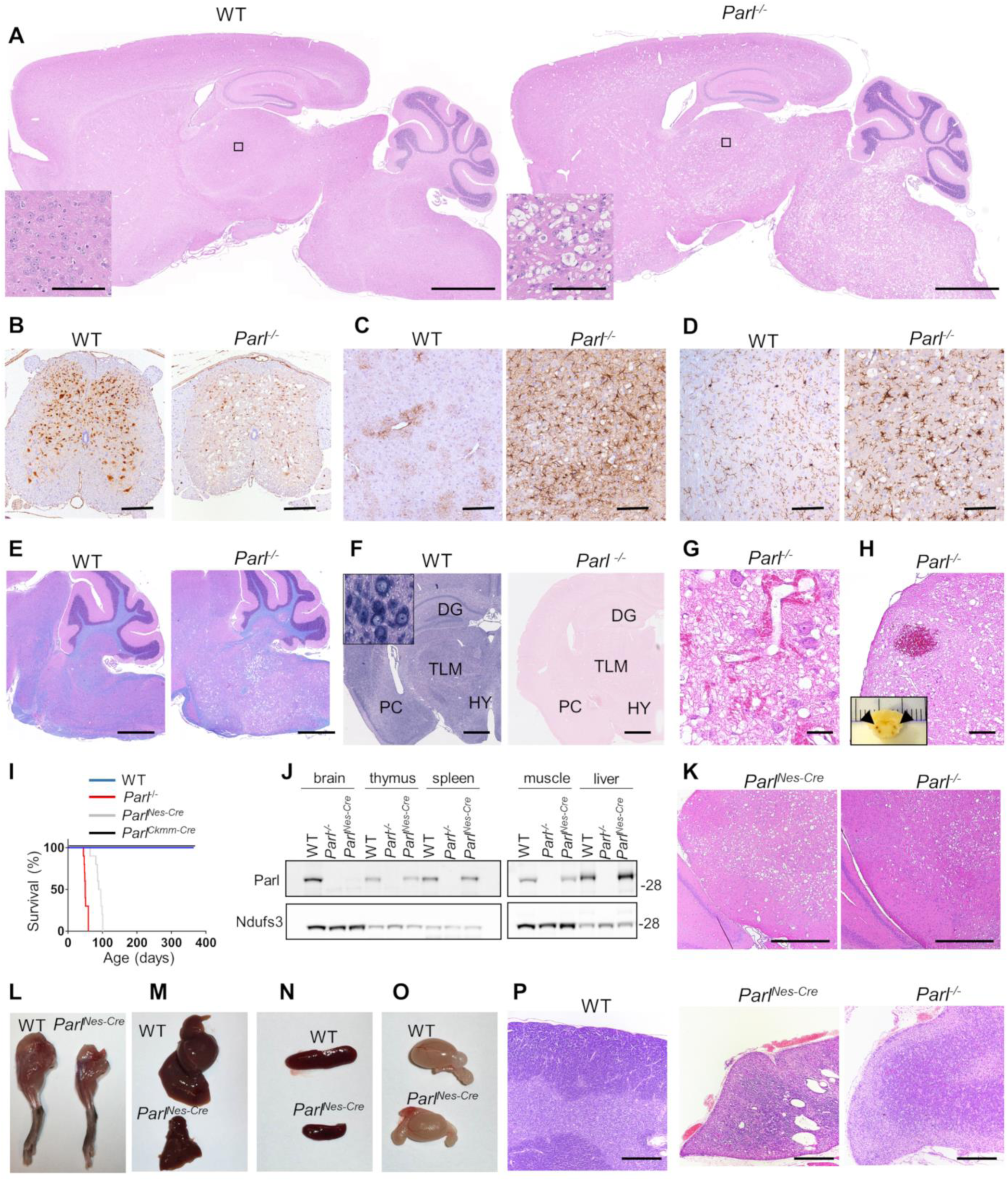
A Leigh-like encephalomyelopathy drives *Parl^-/-^* phenotype. (*A*) Severe vacuolar neurodegeneration in a seven week-old *Parl^-/-^* mouse brain. Representative H&E staining (n≥ 12). Scale bar: 1250 μm. The insert shows a detail corresponding to the thalamus (scale bars: 100 μm). (*B*) Severe neuronal loss in the gray matter of *Parl^-/-^* lumbar spinal cord at 7 weeks of age. NeuN immunoperoxidase staining (n=3 for WT, n=6 for *Parl^-/-^*). Scale bar: 125 μm. (*C*) Anti-Gfap staining showing prominent astrogliosis in *Parl^-/-^* medulla oblongata at 7 weeks of age (n=3 for WT, n=6 for *Parl^-/-^*). Scale bar: 125 μm. (*D*) Anti-Iba1 immunoperoxidase staining in superior colliculus in the midbrain at 7 weeks of age (n=3 for WT, n=6 for *Parl^-/-^*). Scale bar: 125 μm. (*E*) Combined Luxol fast Blue and H&E staining showing preservation of the white matter (stained in blue) in presence of severe neurodegeneration in *Parl^-/-^* mouse brain at 7 weeks of age (n=3 for WT, n=7 for *Parl^-/-^*). Scale bar: 750 μm. (*F*) *Parl* in situ hybridization. DG: dentate gyrus; TLM: thalamus; PC: pyriform cortex; HY: hypothalamus. Scale bar: 1 mm. The insert shows strong Parl expression in WT reticular neurons. (*G*) Representative H&E staining of the inferior colliculus in the midbrain of a 7-week old *Parl^-/-^* mouse showing vascular proliferation (n>10). Scale bar: 50 μm. (*H)* Focal hemorrhage in the olivary nucleus of a 7-week old *Parl^-/-^* mouse. Bilateral symmetrical hemorrhages have been detected in brainstems of 4 out of 7 *Parl^-/-^* mice at 7 weeks of age and of none of the WT littermates. Scale bar: 250 μm. The insert shows that bilateral symmetrical hemorrhagic foci are already visible at gross examination in the medulla oblongata (arrows). (*I*) Kaplan-Meier survival curves of WT (n=14), *Parl^-/-^* (n=14), *Parl^Nes-Cre^* (n=15) and *Parl^Ckmm-Cre^* mice (n=13). (*J*) Representative western blot analysis of Parl protein in brain, thymus, spleen, muscle, and liver mitochondria isolated from 7 weeks old WT and *Parl^-/-^*, and 13 weeks old *Parl^Nes-Cre-^*mice. Ndufs3 is the loading control. (*K*) Representative H&E staining of midbrains from ten weeks old *Parl^Nes-Cre^* (n=4), and seven weeks-old *Parl^-/-^* mice (n=12). Scale bar: 380 μm. (*L-O*) Severe atrophy of the skeletal muscle (*L*), liver (*M*), spleen (*N*) but normal testis size (*O*) in 11-week old *Parl^Nes-Cre-^* male mice compared to an age matched WT littermate control (n>15). (*P*) Representative H&E staining of thymus sections of WT (age: 7 weeks; n=6), *Parl^Nes-Cre^* (age 10-13 weeks; n=4), and *Parl^-/--^* mice (age:7 weeks; n=12) showing severe thymus atrophy. Scale bar: 200 μm.

### Parl deficiency causes early neuronal mitochondrial ultrastructural abnormalities followed by neuronal necrosis

To characterize mitochondrial ultrastructure and the morphological features of cell death induced by Parl deficiency in the brain we performed electron microscopy. Examination of WT and *Parl^-/-^* medulla oblongata at 3 weeks of age, which is before the occurrence of clinical signs or histopathological lesions, shows that *Parl^-/-^* neurons display scattered mitochondria with abnormal cristae (Fig. 2A). At later time points, mitochondrial ultrastructural abnormalities become more severe (Fig. 2B-C) and associate with severe neuronal vacuolization, swelling and loss of integrity (Fig. 2C, 3A-B). This picture is indicative of necrosis, while the typical morphological signs of apoptosis are absent. Similar ultrastructural abnormalities are present in neuronal mitochondria from *Parl^Nes-Cre^* mice (Fig.S1). Since Parl has been linked previously to apoptosis^6,24^, we performed biochemical experiments to investigate further the contribution of this process to the neurodegeneration. During apoptosis, the outer mitochondrial membrane becomes permeable, and cytochrome c is released from the mitochondrial intermembrane space to the cytosol leading to proteolytic activation of executioner caspases and Parp. We analyzed mitochondrial outer membrane permeability of brain mitochondria isolated from symptomatic *Parl^-/-^* mice by measuring the enhancement of complex IV-driven respiration before and after addition of exogenous cytochrome c, which is unable to reach complex IV if outer membranes are intact. Exogenous cytochrome c does not significantly enhance complex IV-driven respiration in both WT and *Parl^-/-^* brain mitochondria (Fig. 3*C*), indicating intactness of outer mitochondrial membranes even in advanced stages of the disease associated with severe neurodegeneration. Consistently, cytochrome c is undetectable in purified brain cytosols from *Parl^-/-^* brains (Fig. 3D), and the expression of full length Parp is similar in *Parl^-/-^* and WT brains without evidence of proteolytic activation (Fig. 3E). Accordingly, we do not see TUNEL positive cells in the degenerating brain areas, notably brainstem, thalamus, hypothalamus at serial time points ranging from asymptomatic to late stage of the disease (Fig. S2A). TUNEL positivity is restricted to few scattered cells, likely undergoing developmental apoptosis, in periventricular areas which are not affected by *Parl^-/-^* neurodegeneration, to a similar extent in WT and *Parl^-/-^* brains. We obtained similar results using antibodies against cleaved caspase 3 (Fig.3G), indicating that apoptosis execution is not overtly impaired in *Parl^-/-^* brains. Conversely, active Caspase 3 immunostaining of atrophic thymus of severely affected *Parl^-/-^* mice shows strong positivity (Fig. S2B) as previously reported ^6^. However, an identical thymus pathology is also seen in late stage *Parl^Nes-Cre^* (Fig. 1P) indicating that this phenotype can be fully induced by deficiency of Parl in the nervous system alone. Treatment of primary *Parl^-/-^* and WT neurons with the pro-apoptotic drug etoposide shows similar proteolytic activation of Caspase 3 and Parp indicating that apoptosis execution is not overtly blocked in cultured neurons (Fig. 3*F*). All together, these data indicate that the striking neurodegeneration induced by Parl deficiency is sustained by neuronal necrosis, without overtly altering the execution of apoptosis in the brain.

**Fig. 2.**
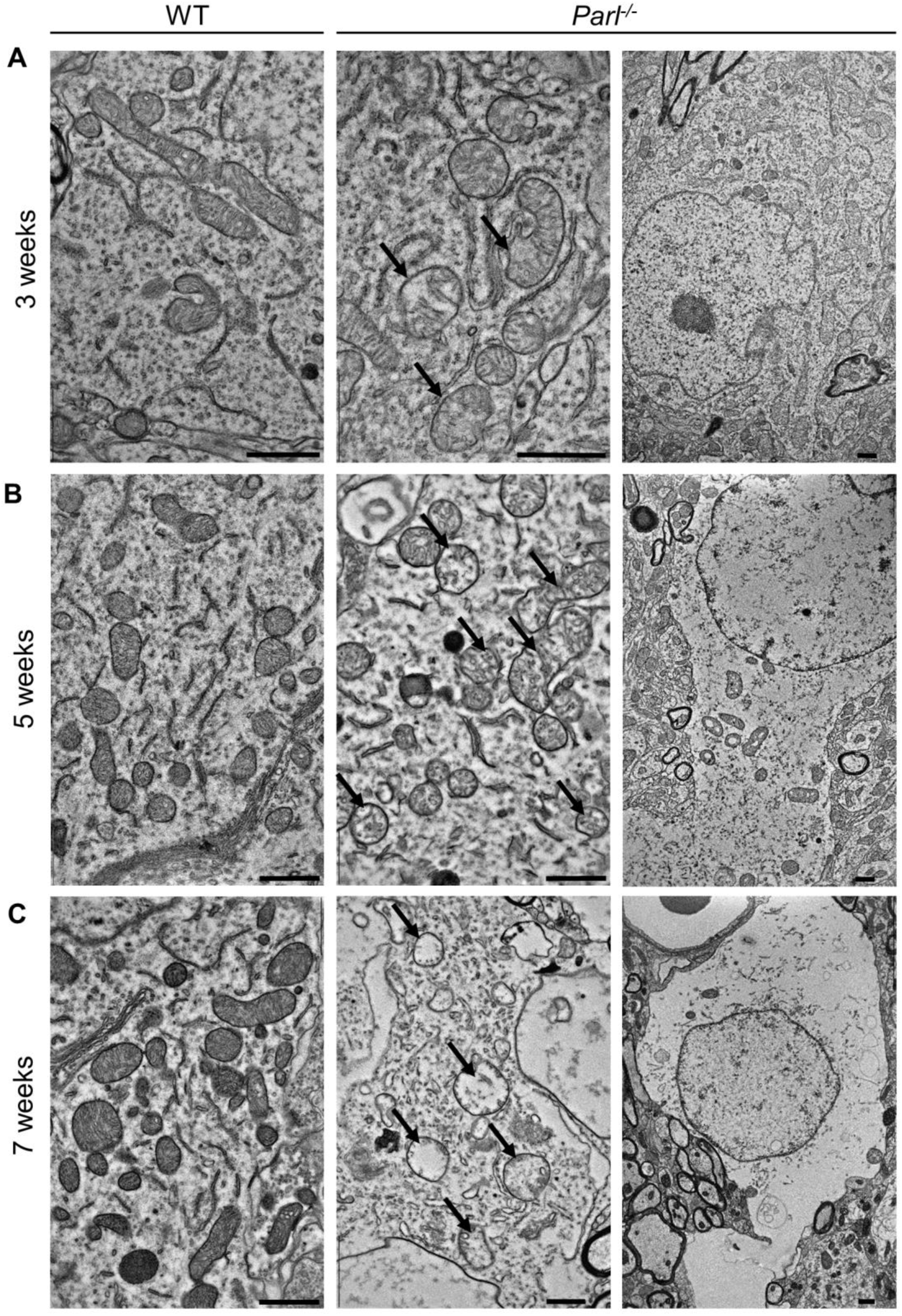
Progressive alterations in *Parl^-/-^* neuronal mitochondrial ultrastructure precede neuronal necrosis. (A) At three weeks of age, transmission electron micrograph shows scattered mitochondria with altered ultrastructure in *Parl^-/-^* neurons of medulla oblongata (black arrows), but not in WT, in absence of necrosis. (B) At the age of five weeks, when necrosis start becoming visible in *Parl^-/-^* brains, mitochondrial structural abnormalities become more diffuse and severe in *Parl^-/-^* neurons (black arrows). (C) At the age of seven weeks worsening mitochondrial morphological abnormalities, characterized by swelling and severe loss of cristae (black arrows), are associated with diffuse neuronal necrosis of *Parl^-/-^* neurons, indicated by cellular swelling, vacuolization and loss of integrity. Scale bar:1 μm.

**Fig. 3.**
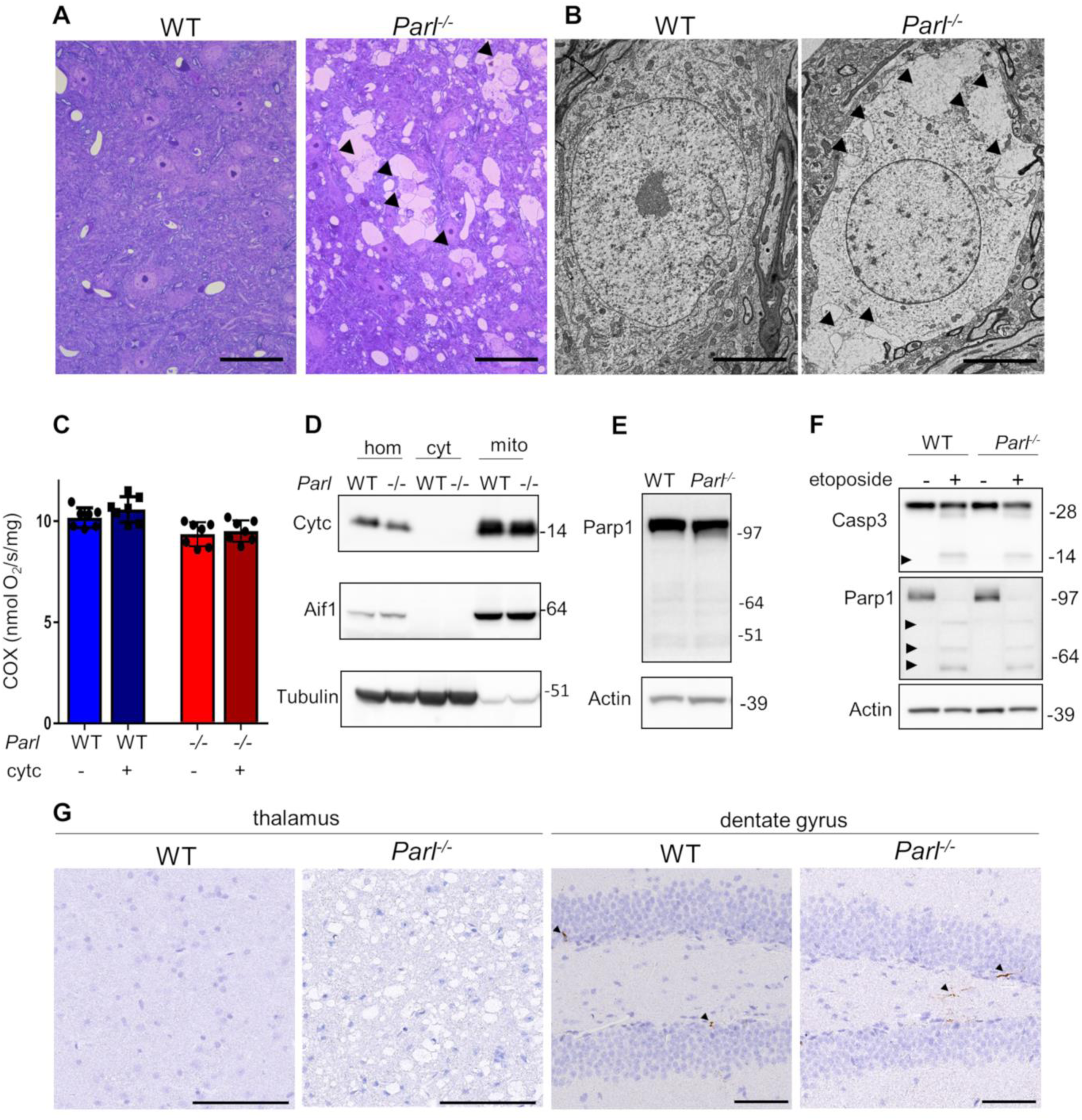
*Parl^-/-^* neurodegeneration is associated with necrotic cell death. (*A*) Semithin section stained with toluidine blue shows vacuolization and disintegration of neurons (black arrowheads) in medulla oblongata of 7 weeks old *Parl^-/-^* mouse compared to a WT littermate. Scale bar: 50μm. (*B*) Representative transmission electron microscopy images showing intracellular vacuolization and loss of membrane integrity in a seven week-old *Parl^-/-^* thalamic neuron. Scale bar: 5μm. (*C*) Evaluation of brain mitochondrial outer membrane permeabilization. Ascorbate/TMPD-driven oxygen consumption rates in 6 weeks old WT and *Parl^-/-^* brain free mitochondria (n=7 for each genotype) before and after addition of exogenous 10 μM cytochrome c. Data represent average ± SD. (*D*) Representative western blot of WT and *Parl^-/-^* brain cytosolic and mitochondrial fractions at 7 weeks of age (n=3 for each genotype) showing absence of cytochrome c into the cytosol. 20 μg of proteins for each fraction were analyzed by SDS-PAGE/immunoblotting using anti-Cytochrome c antibody. Anti-Aif1 and anti-Tubulin are mitochondrial and cytosolic markers respectively. Hom: total homogenate; cyt: purified cytosol; mito: purified mitochondria. (*E*) 20 μg of 7 week-old WT and *Parl^-/-^* nuclei-enriched brain fractions were analyzed by SDS-PAGE/immunoblotting using an anti-Parp antibody. (*F*) Cultured primary neurons were treated with 10 μM etoposide for 24 hours and lysed. 20 μg of total neuronal lysates were analyzed by SDS-PAGE/immunoblotting with anti-Caspase-3 and anti-Parp antibodies. Actin is the loading control. (*G*) Activated caspase 3 staining shows absence of positive neurons, in seven weeks *Parl^-/-^* thalamus despite the presence of severe neurodegeneration. Scale bar: 100 μm. In dentate gyrus, a brain area never affected by *Parl^-/-^* neurodegeneration, activated caspase 3 staining shows scattered positive neurons to the same extent in WT and *Parl^-/-^* mice. Scale bar: 50 μm.

### *Parl^-/-^* Leigh-like encephalomyelopathy is not caused by misprocessing of Pink1 and Pgam5

Next we wondered to which extent the dramatic neurodegenerative phenotype of *Parl^-/-^* mice can be attributed to misprocessing of the best characterized substrates of Parl, Pink^15,10,12^, a mitochondrial kinase, and Pgam5, a mitochondrial phosphatase^13^. Both have roles in neurological diseases^15,16^ and in mitophagy^16,25^. Pgam5 have also been linked to regulation of multiple cell death pathways including necroptosis^26^. However, the contribution of Pink1 and Pgam5 misprocessing to the *Parl^-/-^* phenotype is not known^2^. Pink1 is barely detectable in WT mitochondria, while a remarkable accumulation of Pink1 substrate is seen in *Parl^-/-^* brain mitochondria (Fig. 4A). Similarly, the unprocessed full-length form of Pgam5 strongly accumulates in *Parl^-/-^* mitochondria while the processed Pgam5 forms runs slightly faster than in WT mitochondria, indicating an alternative Parl-independent cleavage. To test whether accumulation of these unprocessed forms of Pink1 and Pgam5 in *Parl^-/-^* mitochondria drives *Parl^-/-^* neurodegeneration, we generated a series of double *Parl^-/-^*/*Pink1^-/-^*, *Parl^-/-^*/*Pgam5^-/-^*, and triple *Parl^-/-^*/*Pink1^-/-^*/*Pgam5^-/-^* combined knock out mice (Fig.4A). Surprisingly, the *Parl^-/-^* phenotype is unmodified by simultaneous deletion of Pink1, or Pgam5, either alone or together, and all these mouse strains invariably died at a similar age as the single *Parl^-/-^* (Fig.4B) affected by similar Leigh like syndrome (Fig.4*C*). To test whether deficient proteolytic products of Pink1 and Pgam5 generated by Parl were essential, we also generated *Pink1^-/-^*/*Pgam5^-/-^*mice. Conversely, *Pink1^-/-^*/*Pgam5^-/-^* mice have a normal lifespan (Fig. 4B) beyond the age of two years without any overt clinical or neuropathological phenotype (Fig. 4C) indicating that misprocessing of Pink1 and Pgam5 alone or together do not explain the *Parl^-/-^* associated Leigh like syndrome

**Fig. 4.**
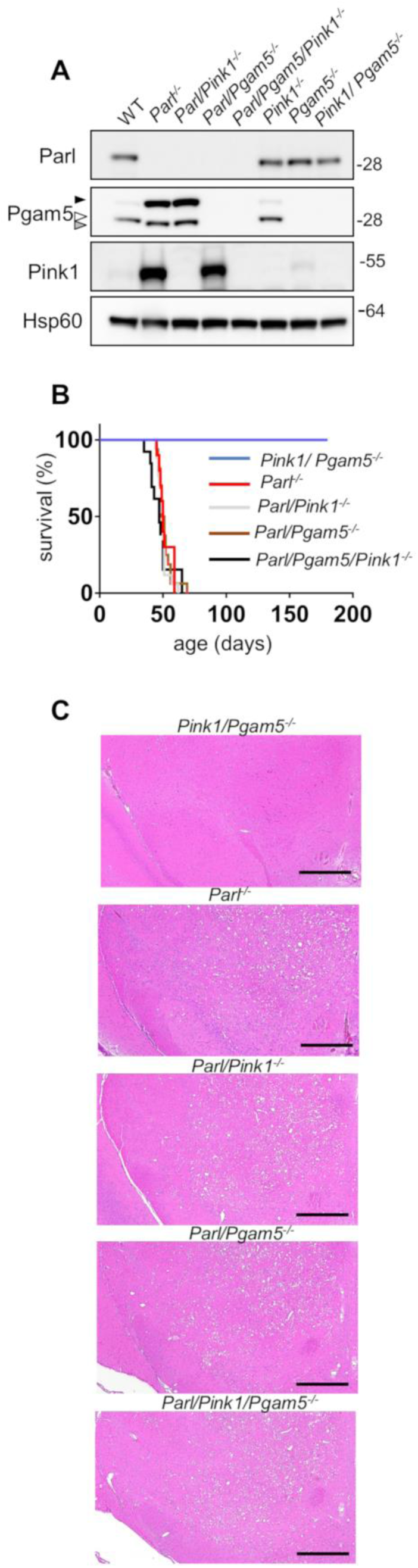
Pink1 and Pgam5 do not interact genetically with Parl deficiency *in vivo*. (*A*) Validation of *Parl*/*Pgam5^-/-^, Parl*/*Pink1*^-/-^ and *Pink1*/*Pgam5*^-/-^ double knockout mice, and *Parl*/*Pgam5/Pink1*^-/-^ triple knockout mice. Brain mitochondria were isolated from mice of the indicated genotype and analysed by SDS page followed by immunoblotting with Parl, Pink1, and Pgam5 antibodies. The white arrow indicates the mature form of Pgam5, the black arrow indicates the unprocessed form, the grey arrow indicates alternatively processed form in *Parl^-/-^* mitochondria. Hsp60 is the loading control. (*B*) Kaplan-Meier survival curves of *Parl^-/-^* (n=10), *Pink1*/*Pgam5^-/-^* (n=10)^-^, *Parl*/*Pink1^-/-^* (n=17), *Parl*/*Pgam5^-/-^* (n=16), *Parl*/*Pink1*/*Pgam5^-/-^* (n=13). (*C*) Representative H&E staining of midbrain coronal sections of seven week-old mice of the indicated genotypes (n>3 for each genotype). Scale bar: 500 μm.

### Mitochondrial functional defects in *Parl^-/-^* brain mitochondria converge on Complex III and Coenzyme Q

Leigh syndrome is a genetically heterogeneous disease caused by different genetic defects that ultimately impair mitochondrial energy production^22^ Therefore, we performed a thorough biochemical characterization of *Parl^-/-^* brain mitochondria to address how mitochondrial function is compromised by absent Parl expression. Since Parl has been previously linked to differences in mitochondrial biogenesis^18^, we asked whether mitochondrial mass is affected in *Parl^-/-^* brains. Expression of respiratory chain subunits and of the outer membrane resident proteins Tomm20 and Vdac1 is similar in WT and *Parl^-/-^* brains at any age (Fig. 5A), indicating unaltered mitochondrial mass. Mitochondrial DNA abundance was also not significantly different between WT and *Parl^-/-^* brains (Fig. 5B). Next, we measured oxygen consumption rates in neuronal mitochondria derived from permeabilized synaptosomes by high-resolution respirometry supplying consecutively substrates and specific inhibitors for Complexes I, II, and IV according to the protocol illustrated in Fig. 5*C*. Importantly, at three weeks of age, which is three weeks before the first clinical signs of *Parl^-/-^* mice, mitochondrial respiration is comparable between WT and *Parl^-/-^* brain mitochondria (Fig. S3A). However, at six weeks of age, both ADP stimulated respiration (OXPHOS), providing a measurement of the maximal capacity to generate ATP, and uncoupled respiration, providing an estimate of the maximal electron transfer capacity (ET), are severely diminished in *Parl^-/-^* neuronal mitochondria (Fig. 5D). This was similar whether they were supplied with only Complex I substrates (CI OXPHOS), or by combined supply of Complex I and II (CI+II OXPHOS and ET). Cytochrome c oxidase, catalyzing the final step in the electron transfer chain, is only slightly, although still significantly compromised (Fig. 5D). Thus, the mitochondrial respiration defect is limited by impaired electron transfer capacity particularly in respiratory chain segments proximal to cytochrome c oxidase. Next, to localize precisely the respiratory chain defect, we measured the maximal enzymatic activities of each of the respiratory chain complexes (CI-CIV) and of ATP synthase in brain mitochondria, as well as the coupled enzymatic activity of CII+III. This latter enzymatic assay explores the capacity of mitochondria supplied with succinate to transfer electrons to cytochrome c, therefore exploring the overall integrity of the respiratory chain in the segment Complex II-coenzyme Q-Complex III. We detected a severe enzymatic defect of Complex III and of Complex II+III (Fig. 5*E*), while the enzymatic activities of Complex II and IV are slightly, although significantly, decreased. Next, we wondered whether the deficient Complex III activity was due to abnormal Complex III assembly. Blue native gel electrophoresis of brain mitochondria showed an effective assembly and maturation of the four respiratory chain complexes I-IV and of ATP synthase, as well as preserved formation of the super complex (Fig. 5F). The terminal component of complex III, the iron sulfur cluster protein Uqcrfs1, is normally incorporated ^27^ but complex III showed consistently a slightly lower electrophoretic mobility in *Parl^-/-^* mitochondria (Fig. 5F). An identical CIII_2_ migration abnormality has been recently reported in mitochondria deficient of Ttc19^28^, a protein required for the maintenance of Complex III enzymatic activity through the regulation of Uqcrfs1 turnover. Interestingly, gene mutations in *TTC19* cause human Leigh syndrome^29^. Next to investigate unambiguously the possibility of a CoQ deficiency in presence of a simultaneous enzymatic defect of isolated Complex III and CII+III (Fig.5E), we directly measured CoQ levels (CoQ_9+10_) in whole brain extracts and the ratio between reduced and oxidized CoQ, a sensitive marker of the electron transfer efficiency of the respiratory chain^30^. CoQ deficiency is one of the established causes of Leigh syndrome^22^. Remarkably, total CoQ levels are severely decreased in *Parl^-/-^* brains at seven weeks of age (Fig. 5G). Moreover, the ratios between reduced and oxidized forms of both forms of CoQ are dramatically increased (Fig. 5H) indicating a marked impairment in CoQH_2_ oxidation, consistent with the impaired Complex III activity identified above. These changes are, in contrast to the respiration defect, already severe at three weeks of age (Fig.S3B) when the mice are asymptomatic, indicating that CoQ deficiency is an early effect of Parl deficiency. In conclusion, Parl is required to prevent severe mitochondrial dysfunction in brain mitochondria characterized by Complex III and CoQ biosynthesis defects.

**Fig. 5.**
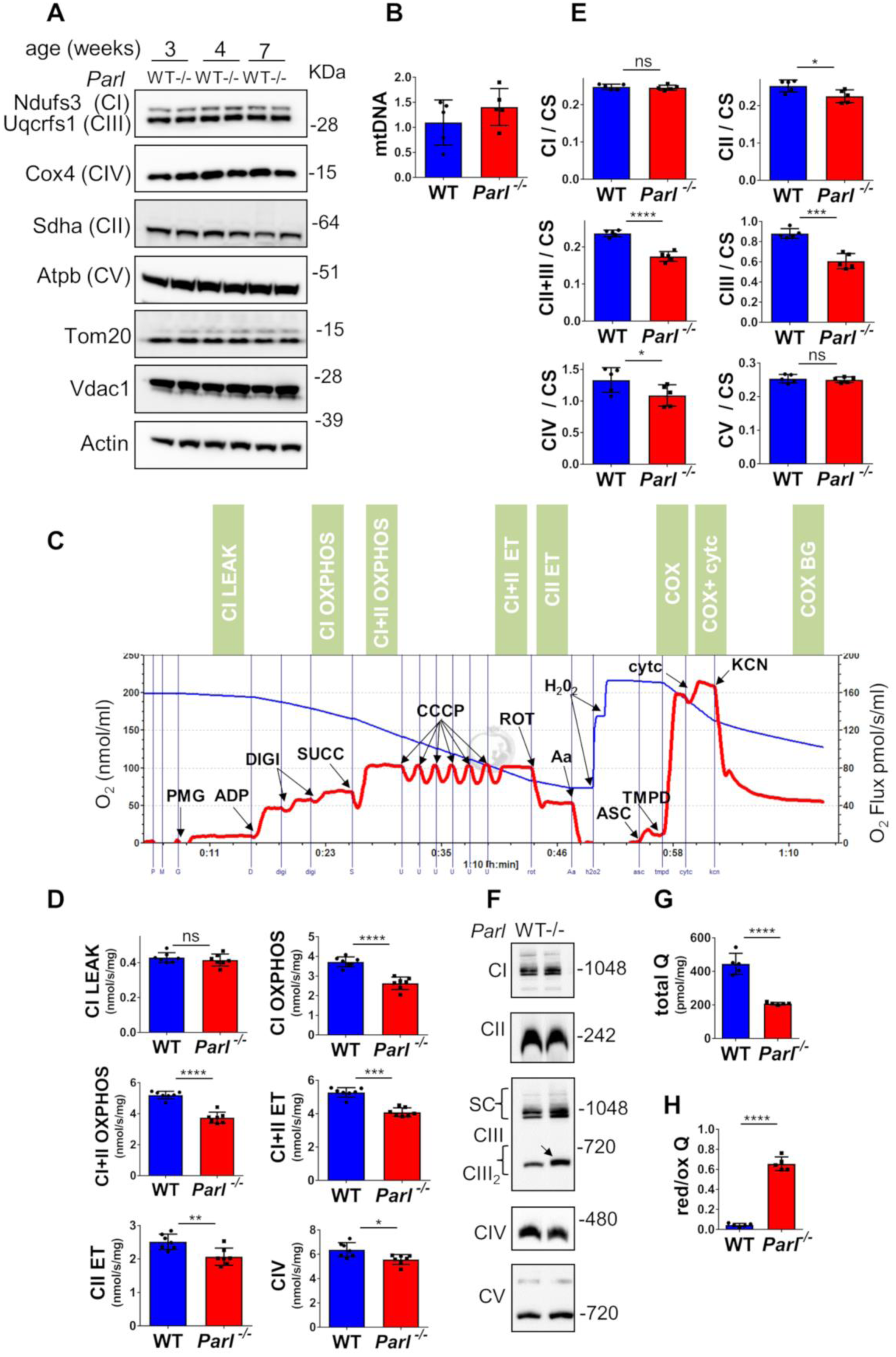
Defects in complex III and Coenzyme Q in *Parl^-/-^* brain mitochondria. (*A*) Representative immunoblot of mitochondrial respiratory chain subunits (CI-CV), Tom20, and Vdac1 in WT and *Parl^-/-^* whole brain lysates at 3, 4, and 7 weeks of age (n=4). Actin is the loading control. (*B*) Mitochondrial DNA, normalized by nuclear DNA, was measured in 7 weeks old WT and *Parl^-/-^* brainstems as specified in the methods section. (*C*) Representative trace illustrating the protocol for high-resolution respirometry in neuronal mitochondria. The blue trace indicates the O2 concentration, the red trace indicates its time derivative. 50 μg purified synaptosomes are loaded in the Oroboros 2k chamber in Miro6 buffer at 37 C. Digitonin (Digi) is titrated to achieve optimal synaptosomal permeabilization. Substrates: Complex I (PMG=pyruvate+malate+glutamate), Complex II (Succ=succinate); Complex IV (ASC/TMPD=ascorbate+TMPD). Uncoupler: CCCP. Inhibitors: Complex I (ROT: rotenone); Complex III (Aa: Antimycin a); Complex IV (KCN: potassium cyanide). Cyt c: cytochrome c. Respiratory states are indicated in the green boxes. CI LEAK=Complex I-driven leak respiration. CI OXPHOS: Complex I-driven phosphorylating respiration. CI+II OXPHOS: phosphorylating respiration driven by simultaneous activation of Complex I and II. CI+II ET: electron transfer capacity driven by simultaneous activation of Complex I and II. CII ET: electron transfer capacity driven by Complex II. CIV: cyanide sensitive cytochrome c oxidase-driven respiration. Cytc: exogenous cytochrome c is added to evaluate the integrity of the outer mitochondrial membranes. H_2_O_2_ in the presence of catalase is used to reoxygenate the chamber when needed. (*D*) Quantification of the respiratory states of permeabilized synaptosomes isolated from six-weeks old WT and *Parl^-/-^* mice (n=7 for each genotype) as from the protocol in C. (*E*) Enzymatic activities of individual respiratory chain complexes and CII+III in brain mitochondria obtained from six weeks old WT and *Parl^-/-^* mice (n=5 for each genotype) by spectrophotometry, and normalized to citrate synthase activity. (*F*) Blue native gel electrophoresis of purified brain mitochondria from 7 weeks old WT and *Parl^-/-^* mice, followed by immunoblotting with anti-Ndufs3 (Complex I), anti-Sdha (Complex II), anti-Uqcrfs1 (Complex III), anti-Cox4 (Complex IV), anti-Atpb (Complex V). The arrow indicates the upward mobility change of dimeric Complex III in *Parl^-/-^*. (*G*) Concentration of total CoQ (Q_9_+Q_10_) measured by HPLC in brain tissue extracts from seven weeks old WT and *Parl^-/-^* mice (n=5 for each genotype).(*H*) Ratio between reduced and oxidized CoQ from the experiment in G. The bar graphs indicate the average ± SD. Statistical significance has been calculated by two sided student t test: *= p<0,05; **=p<0,01; ***=p<0,001;****=p<0,0001.

### Mitochondrial proteome changes induced by Parl deficieny in the brain

We wondered what mitochondrial protein changes underlie the*Parl^-/-^* Leigh-like phenotype and the observed Complex III and CoQ biosynthesis defects. We performed a mass spectrometry-based proteome analysis of brain mitochondria purified from three WT and three *Parl^-/-^* mice, leading to the quantification of 781 out of 1085 proteins annotated in the mouse mitochondrial proteome in Swissprot. The volcano plot shows that despite the dramatic phenotype, surprisingly few mitochondrial proteins are differentially expressed in *Parl^-/-^* compared to WT brains (Fig. 6A), indicating selective effects of Parl on the mitochondrial proteome. Among these, we find a striking downregulation of the Complex III regulating protein Ttc19^28,31^ and of several proteins required for CoQ biosynthesis (Coq3, Coq4, Coq5). We also notice the significant decrease of the sulfide-CoQ oxidoreductase Sqrdl, which has been recently reported to decrease in human diseases caused by primary CoQ defects^32^. Interestingly, other known CoQ oxidoreductases such as electron transfer flavoprotein-ubiquinone oxidoreductase (Etfdh), glycerol 3-phosphate dehydrogenase (Gpd2), dihydroorotate dehydrogenase (Dhod), proline-dehydrogenase (Prodh), and choline dehydrogenase (Chdh) are not affected. Ghitm, a multipass inner membrane protein not previously linked to Parl, also decreases. Conversely, similar Pgam5 also Diablo14 is increased in Parl deficient brain mitochondria (Fig. 6B). Next, we validated these findings by western blot in brain mitochondria from WT, *Parl^-/-^*, and *Parl^Nes-Cre^* (Fig. 6*B*). Since our proteomic approach identifies substrates by expression changes, we decided to include in the validation also the previously reported Parl substrates Htra2^9^, Opa1^6^, Stard7, and Clpb^14^, which were not significantly different in our mass spectrometry analysis (Fig. 6A). Western blots confirm the virtual disappearance of Ttc19, a marked reduction of the CoQ proteins (Coq3,Coq4,Coq5), Sqrdl, and Ghitm, and altered cleavage of Ttc19, Pgam5, Diablo, Stard7, and Clpb. The expression and processing of Htra2 and Opa1 are not impaired (Fig. 6B). To correlate these protein changes with the clinical phenotype, we checked their expression in brain tissue over time, at the age of 1, 3, and 5 weeks, when *Parl^-/-^* mice are still asymptomatic, and at 7 weeks, when the mice are severely affected by neurological deficits (Fig. 6C, Fig. S4). To further investigate the molecular basis of the Complex III and CoQ deficiency of *Parl^-/-^* brains we included in this time course an extensive panel of proteins required for CoQ biosynthesis. In the absence of Parl, we notice the virtual disappearance of mature Ttc19 at all ages indicating that Parl is required for Ttc19 maturation and expression, and a severe reduction of Coq4 already noticeable at 1 week of age (Fig. 6C), followed by decrease in Coq3, Coq5, Coq6, Coq7, Coq9 and of Sqrdl at later time points.

**Fig. 6.**
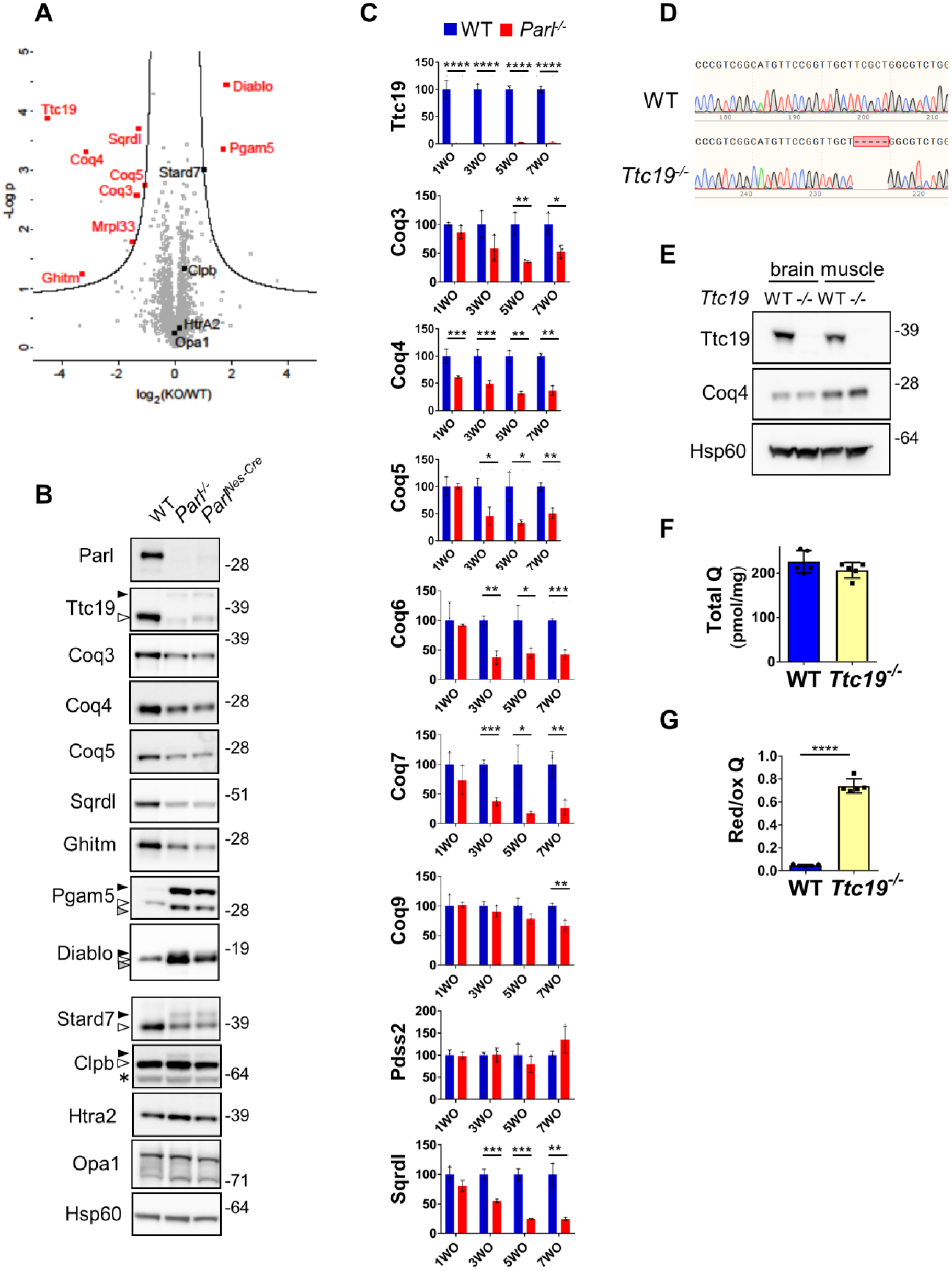
Restricted changes in the brain mitochondrial proteome induced by Parl deficiency explain the Complex III and CoQ defects (*A*) Volcano plot showing differentially regulated proteins in *Parl^-/-^* brain mitochondria. purified from 5 week-old WT and *Parl^-/-^* brains (n=3) analyzed by mass spectrometry. Significantly differentially regulated proteins are distributed outside the volcano cutoff of fold changes >2 and p value <0.05. Significantly differentially expressed mitochondrial proteins are plotted in red. Previously reported Parl substrates that did not reach statistical significance are plotted in black. The non-mitochondrial proteins Trabd, Bcap31, Bcan, Eif1a, and Setdb, appeared similarly expressed as from the validation experiment in Fig.5B and therefore were not included in the graph. (*B*) Validation of the mass spectrometry results and of previously reported Parl substrates Stard7, Clpb, Htra2, Opa1. Twenty μg of mitochondria isolated from brain of WT, *Parl^-/-^*, and *Parl^Nes-Cre^*, were analyzed by SDS page followed by immunoblotting. The white arrows indicate the mature (processed) form of the protein, the black arrows indicate unprocessed forms, the grey arrows indicate alternatively processed forms in *Parl^-/-^* mitochondria. The asterisk indicates bands of uncertain significance. (*C*) Time course of the protein expression of Ttc19, CoQ biosynthesis proteins and Sqrdl in WT and *Parl^-/-^* brains. 20 μg of total brain lysates from WT, *Parl^-/-^* sacrificed at 1, 3, 5, and 7 weeks of age (n=3), were analyzed by SDS page followed by immunoblotting. The graph bars indicate the quantifications of the western blot results, normalized with the loading control Hsp60 and expressed as % relative to the WT. The original western blots are shown in *SI Appendix*, Fig.S5. (*D*) Generation of *Ttc19^-/-^* mice by Crisper/Cas9 technology. *Ttc19^-/-^* mice have a 5 base pairs deletion in the first exon. (*E*) Immunoblot analysis of brain and muscle mitochondria (20 μg) with anti Ttc19 and anti Coq4 antibodies. Hsp60 is the loading control. (*F*) Concentration of total CoQ Q (Q_9_+Q_10_) were measured by HPLC in brain tissue from7 weeks old WT and *Ttc19^-/-^* mice (n=5 for each genotype). (*G*) Ratio between reduced and oxidized CoQ from 7 weeks old WT and *Ttc19^-/-^* mice from the experiment in F. The bar graphs indicate the average ± SD. Statistical significance has been calculated by two sided student t test: ****=p<0,0001.

To explore to what extent the catalytic function of Parl is involved in these protein changes, we checked the expression of these proteins in stable *Parl^-/-^* MEFs expressing Parl^WT^ or catalytic inactive Parl^S275A^ (Fig. S5). We observed no altered mobility of Coq4, Coq5, Sqrdl in *Parl^-/-^* cultured MEFs, suggesting that they likely are not direct Parl substrates. In contrast, the processing of Ttc19, Pink1, Pgam5, Stard7, Diablo, and Clpb is clearly modified by expression of catalytically active Parl, but not by mutant Parl^S275A^, confirming that Parl proteolytic activity is required for the maturation of these proteins and that they are all likely genuine Parl substrates^14^. In contrast, we could again not confirm differences in Htra2 and Opa1 processing in *Parl^-/-^* cultured MEFs suggesting that both these proteins are not Parl substrates in contrast to what previously proposed^6,9^.

To dissect the relationship between the Ttc19 deficiency and the observed CoQ defects we generated *Ttc19^-/-^* mice (Fig. 6D-E). These mice (20 out of 20) have not shown any reduced survival although they seem to show a slight reduction in locomotor activity at one year of age. Coq4 expression is normal in *Ttc19^-/-^* (Fig. 6E) as well as CoQ concentration (Fig. 6F). Conversely, CoQ red/ox ratio is dramatically increased in *Ttc19^-/-^* brains (Fig. 6G), as in the *Parl^-/-^* (Fig. 5*G*). Thus, deficiency of mature Ttc19 in *Parl^-/-^* brains explains the abnormally increased CoQ red/ox ratio, consistent with the defective Complex III activity, but not the CoQ biosynthesis deficit. Altogether, the data indicate that Parl is required for the maintenance of Complex III activity by stabilization of Ttc19, and for efficient CoQ biosynthesis by stabilizing Coq4 expression (Fig. 7A-B).

**Fig. 7.**
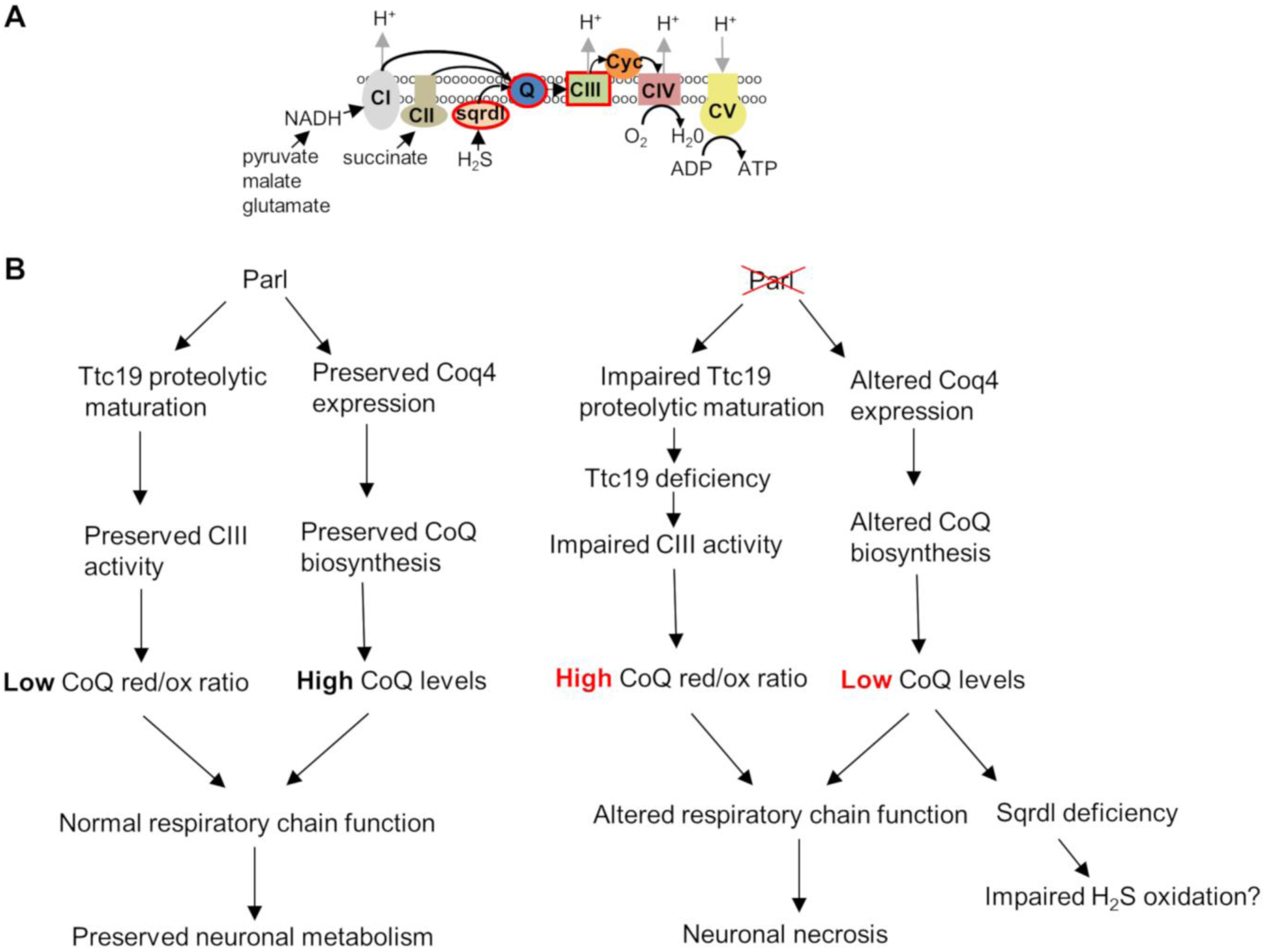
Schematic illustration of the role of Parl in neuronal mitochondria homeostasis. (*A*) Cartoon illustrating the mitochondrial respiratory chain. CI: Complex I; CII: Complex II; CIII: Complex III; CIV: Complex IV; CV: ATP synthase; Sqrdl:sulfide quinone oxidoreductase; Q: Coenzyme Q; cyc: cytochrome c. The sites encircles in red are affected by Parl deficiency in neuronal mitochondria. Coenzyme Q, a lipophilic quinone embedded in the inner mitochondrial membrane, constitutes an electron transfer chain relay site, receiving electrons from Complex I (derived from NADH oxidation), from complex II (derived from succinate oxidation), from Sqrdl (derived from sulfide oxidation) and from other metabolic enzymes. Electrons are then transferred from reduced coenzyme Q to cytochrome c by complex III, then from cytochrome c to molecular oxygen by complex IV. (*B*) Schematic diagram illustrating how Parl expression regulates the respiratory chain function in the nervous system.

## DISCUSSION

This work reveals an essential role of Parl for the homeostasis of the nervous system. Deficiency of Parl in the nervous tissue whether isolated or in the context of systemic Parl knockout, leads to a striking neurodegeneration rather similar, from a neuropathological point of view, to the phenotype of the Complex I deficient *Ndufs4^-/-^* mouse^33,34^, the only currently available mouse model for Leigh syndrome. Although much caution is needed when extrapolating from mouse models to human diseases, this study prompts to investigate whether or not *PARL* gene mutations are present in human patients with unexplained Leigh disease or Leigh-like syndromes. Deletion of Parl in the nervous system alone in *Parl^Nes-Cre^* largely mimics the phenotype of germline *Parl^-/-^*, as *Ndufs4^Nes-Cre^* recapitulates that of germline *Ndufs4^-/-^*, indicating that in both models the nervous system involvement drives the disease. The molecular basis for this remarkable tissue specificity remains unclear for mouse and human Leigh syndrome, as well as for most of the major neurological diseases. Nevertheless, this observation demonstrates that the Leigh syndrome is actually the elusive cause of the multisystem lethal phenotype of the previously generated *Parl^-/-^* mice^6^. Interestingly, severe neurological alterations have also been observed in flies carrying mutations in the Parl fly orthologue rhomboid-7^35^.

Previous work focused on the potential role of Parl in apoptosis but has not noticed this severe neuropathological phenotype^6^. We are currently investigating why this phenotype was overlooked before. From the current data it is in any event clear that the specific deletion of Parl in the nervous system alone in the *Parl^Nes-Cre^* mice is sufficient to cause a similar lethal multisystem phenotype as germline *Parl^-/-^* mice, including severe atrophy of skeletal muscle thymus and spleen atrophy^6^, although with a delay of about one month. Theoretically, we cannot fully exclude nonspecific Cre recombinase activity driven by Nestin in scattered cells in tissues outside the nervous system (https://www.jax.org/strain/003771). However, as shown in Fig 1J Parl protein expression in these tissues was consistently comparable to WT animals suggesting that nonspecific deletion of Parl in the peripheral tissues of *Parl^Nes-Cre^* mice is minimal. Moreover, the lack of overt muscle atrophy and lethality in mice with specific deletion of Parl in muscle, as opposed to the nervous system, indicates that the severe muscle atrophy is mainly caused by the neurodegeneration. This might be due to denervation, and this hypothesis is supported by the widespread degeneration of neurons in the spinal cord grey matter, including in the anterior horns where cell bodies of the lower motor neurons are localized. Brain specific deletion of the mitochondrial protease Htra2 affects thymus and spleen in a similar way as brain specific Parl*-* deficiency^36^. Such immunological phenotypes can be caused by a variety of general stress conditions that trigger increased secretion of corticosteroids and other stress response hormones^37^. Hind limb unloading for instance induces the distinctive depletion of double positive CD4+CD8+ T lymphocytes that is also observed in *Parl^-/-^* thymus and drop of B lymphocytes in *Parl^-/-^* spleen^6^, but other general stress conditions such as malnutrition^38^, amyotrophic lateral sclerosis^39^ and cerebrovascular accidents^40^ can have similar effects on thymus and spleen. This is consistent with the observation that *Parl^-/-^* and *Parl^Nes-Cre^* mice develop these immunological manifestations only in late stages of the disease when they are already affected by severe neurological deterioration.

Surprisingly, despite the striking neuropathological phenotype, the absence of Parl affects only a very circumscribed proportion of the brain mitochondrial proteome, which overlaps with recent proteomic data obtained in HEK293 cells by Chafradic and quantitative affinity purification^14^. At odd with previous observations^6,9^ and consistent with following reports^14,41^, the data presented here do not support Opa1 and Htra2 as substrates of Parl.

Conversely, we find that several of the few proteins that are differentially regulated by Parl are known to play important roles in neurological diseases. We tested explicitly whether the accumulation of Pink1 and Pgam5, two substrates of Parl known to be involved in Parkinson’s disease^15,16^ and mitophagy^16,25^, could be responsible for the pathological phenotype of the *Parl-/-* mice. Pgam5 has also been reported to modulate necroptosis and other cell death modalities at least *in vitro*^26^. Rare *PARL* mutations have been reported in few patients with Parkinsons’s disease^5^. We tested combined knockouts of these two substrates with *Parl^-/-^* to evaluate whether their accumulation caused the lethality. We also tested separately their combined deletion in the presence of Parl to check whether the loss of function of these two substrates would play a role in the *Parl^-/-^* phenotype. Both experiments did not modulate or simulate the *Parl^-/-^* phenotype. Thus Pink1 and Pgam^515,16,25^ are not responsible for the Leigh-like pathology in the *Parl^-/-^* mice suggesting that other biochemical mechanisms are involved. Moreover, the normal survival and lack of striking clinical phenotypes in *Pink1^-/-^*/*Pgam5^-/-^* suggests that these proteins are not essential for mitochondrial turnover *in vivo*.

Here we show that Parl is essential for the maintenance of the respiratory chain function independently from effects on mitochondrial mass^18^ by regulating Complex III activity and CoQ biosynthesis. Parl is required for the expression of Ttc19 (Fig. 6*B-C* and Saita et al.^14^). Impaired proteolytic maturation of Ttc19 by Parl likely leads to its degradation by alternative mitochondrial proteases. Ttc19 is involved in the turnover of the iron-sulfur protein Uqcrfs1^28^, a structural subunit of Complex III essential for the catalytic activity that ensures CoQ oxidation. When this activity is normal, CoQ red/ox ratio is low, while it is high when Complex III is dysfunctional, as in the case of the *Parl^-/-^* mice. In addition, and independently from Ttc19, Parl ensures CoQ biosynthesis by stabilization of Coq4, a protein required for the biosynthesis of this electron acceptor^42^. Intriguingly, the profound CoQ deficiency that we found in *Parl^-/-^* brains is followed by a secondary reduction in the expression of Sqrdl, as also recently observed in models of primary CoQ defects^32,43^. This hydrophobic molecule serves essential functions in mitochondrial membranes both as an electron carrier from several metabolic pathways to Complex III and as membrane antioxidant^44,45^. The mechanism that links ablation of Parl with the reduced Coq4 and the resulting CoQ deficiency is however not yet clear, since we could not demonstrate that Coq4 is a substrate of Parl. One possibility is that misprocessing of a still undefined Parl substrate affects CoQ biosynthesis upstream of Coq4, similarly to how the yeast intermediate peptidase Oct1p ensures CoQ biosynthesis by cleaving Coq5p^46^. An alternative hypothesis is that Parl regulates CoQ biosynthesis indirectly. Recent observations show that the CoQ pathway is sensitive to different molecular mechanisms affecting mitochondrial DNA gene expression in mouse heart tissue^47^. However, in these models Smac and Parl expression was upregulated, while Coq4 expression was unmodified or increased, in contrast with what we observe in *Parl^-/-^* brains, where additionally the expression of mitochondrial DNA encoded proteins is unaffected. Thus, the effects caused by Parl deficiency are clearly different from those reported in this study. Similarly, it remains unclear how Parl consistently affects the expression of the inner membrane protein Ghitm, which appears to be consistently downregulated in absence of Parl in different cell types without obvious evidence of being a Parl substrate, raising the possibility that the effect of Parl on Coq4 and Ghitm expression might be also independent from its proteolytic activity. The *Ttc19^-/-^* mice data indicate that the CoQ deficiency is independent from the complex III defect. Interestingly, human mutations in *TTC19*^29^, and in genes that induce CoQ deficiency ^22^ are independent causes of Leigh syndrome^48^. In addition, *COQ4* haploinsufficiency is sufficient to cause neurodegeneration in human^48^. Although deletion of Ttc19 only causes mild neurodegeneration in mouse^28^, we propose that the combination of the defects in Complex III activity and in CoQ biosynthesis could play an important role in the severe mitochondrial dysfunction that we observe in *Parl^-/-^* neuronal mitochondria.

Since the original description of *Parl^-/-^* mice^6^, many studies have focused on the possible link between Parl and apoptosis *in vitro*. The results of these investigations have been controversial and both pro- and anti-apoptotic mechanisms have been proposed^6,9,24^. Here we show that Parl deficiency leads to neuronal necrosis *in vivo*, and that the effects on apoptosis *in vivo*, such as those observed in thymus and spleen, appear largely indirect. We speculate that the necrotic phenotype we observed in the nervous system may represent a tissue specific consequence of the severe structural and functional damage that progressively accumulate in neuronal mitochondria, rather than the result of a direct gatekeeping activity of Parl in regulated necrosis machineries. Although we were not able to define in further detail whether necrosis is regulated or not in *Parl^-/-^* brains, the neuropathological phenotype is consistent with the definition of Leigh syndrome as a necrotizing encephalomyelopathy. Future studies may investigate whether any subroutine of regulated necrosis is implicated or not in *Parl^-/-^* neurodegeneration. Our work indicates that Parl has a constitutive physiological role in keeping the respiratory chain function and structural integrity in check, with important consequences for the nervous system.

## MATERIALS AND METHODS

### Mice

Mice conditionally targeted and full knock out for *Parl*tm1Bdes (*Parl^lox/lox^*), *Parl*tm1.1Bdes(*Parl*^-/-^) were previously generated^6^. Nervous system specific *Parl*^-/-^ mice, called *Parl^Nes-Cre^*, were generated by crosses with *Tg(Nes-Cre)1Kln*^49^ mice that express the recombinase Cre in the central and peripheral nervous system and muscle specific *Parl*^-/-^ mice, called *Parl^Ckmm-Cre^*, were generated by crosses with *Tg(Ckmm-Cre)5Khn*^50^. Animals are heterozygous for the *Cre* transgene and littermates not having the *Cre* transgene were used as controls. *Pgam5*tm1d(EUCOMM)Wtsi (*Pgam5^-^*^/-^) full KO mice were obtained after crossing *Pgam5*tm1a(EUCOMM)Wtsi with Gt(ROSA)26Sortm1(*Flp1*)Dym to obtain conditionally targeted *Pgam5*tm1b(EUCOMM)Wtsi mice which were finally crossed with Tg(Ella-*Cre*)C537Lmgd/J. Mice negative for the *Flp* and *Cre* transgenes were used for further breedings. *Pink1*tm1.1Wrst (*Pink1^-/-^*) mice were kindly provide by Wolfgang Wurst ^51^. *Ttc19^-/-^* mice were generated using Criper/Cas9 technology to target the first exon, encoding the mitochondrial sorting motif. mRNA guides were selected using the CRISPOR web tool (http://crispor.tefor.net/) and tested by a T7 endonuclease assay in mouse embryonic stem cells. Guide 5’-GCAUGUUCCGGUUGCUUCGC-3’ was selected. Ribonucleoproteins (RNPs) containing 0.3 μM purified Cas9 protein (Integrated DNA Technologies, IDT) and 0.6 μM CRISPR RNA crRNA and *trans* activating crRNA (IDT) were injected into the pronucleus of C57Bl6J embryos by microinjection in the MutaMouse facility of the KU Leuven. F0 mice were genotyped after weaning using PCR and Sanger sequencing. Founder mice carrying a deletion (D) of 5 nucleotides (D5 +15bp to +19bp, with A of the ATG start codon at position +1) were crossed once with C57Bl/6J and homozygous mice were obtained after breeding heterozygous offspring. The absence of protein or truncated forms was confirmed by western blot analysis using the C-terminal specific Anti-TTC19 antibody anti-ttc19 (Sigma HPA052380). Mice were inbred on a *C57Bl*/*6J* background and both females and males were included in the study. Mice are housed in cages enriched with wood wool and shavings as bedding, and given access to water and food ad libitum. All experiments were approved by the Ethical Committee on Animal Experimentation of the University of Leuven (KU Leuven).

### Pathological analysis

*Parl*^-/-^ mice and matched *WT* controls at 2-8 weeks of age were used. Animals were anesthetized with a mixture of xylazine (25 mg/ml), ketamine (20mg/ml) and atropine (20ng/ml), followed by intracardial perfusion of ice-cold phosphate buffered saline (PBS, 10 mL) and ice-cold paraformaldehyde (4% PFA). Head and spine were fixed for additional 48 hrs in 4% PFA at 4 °C. After fixation, brain was removed from the skull and serially sliced using the coronal brain matrix system (Zivic Instruments BSMYS001-1). Five slices were obtained from each brain using the following anatomical landmarks as references: paraflocculi, mammillary body, optic chiasm, olfactory tubercles. Spine was decalcified in a supersaturated solution of tetrasodium EDTA and cross-sectioned at the level of cervical, thoracic and lumbar segments. Samples were embedded in paraffin blocks. Five μm thick sections were stained with hematoxylin and eosin (H&E). Some sections were stained with Luxol Fast Blue, TUNEL or with antibodies against Gfap, Iba-1, NeuN, or Cleaved Caspase-3. Microscopic lesions were classified according to the International Harmonization of Nomenclature and Diagnostic Criteria for Lesions in Rats and Mice INHAND criteria^52^

### In situ hybridization

In situ hybridization (ISH) was performed as described previously^53^. A cDNA-containing construct of *Parl* was generated in the pGemT-easy vector system (Promega). Digoxigenin-labeled antisense and sense RNA probes were generated by in vitro transcription of linearized plasmid using T7 and SP6 RNA polymerase (Roche), respectively. Brains were collected, fixed with 4% paraformaldehyde overnight, washed with PBS, dehydrated and embedded in paraffin. ISH was performed on 6?μm brain sections using an automated platform (Ventana Discovery, Ventana Medical Systems). Images were obtained by light microscopy and captured with a CH250 CCD camera (Photometrics, Tucson, AZ).

### Electron microscopy

Mice of the indicated genotype were transcardially perfused with 2.5% glutaraldehyde, 2% paraformaldehyde in 0.1 M cacodylate buffer pH 7.4 and their brains were post-fixed in the same solution overnight at 4°C. Coronal brain sections (300 μm thick) were cut on a Leica VT1000S vibratome and rectangular pieces of tissue comprising the regions of interest (thalamus, cortex and medulla oblongata) were dissected. The tissue was stored overnight at 4°C in the fixative solution, washed in 0.1 M cacodylate buffer and post-fixed for 2 hours at RT with 1% OsO_4_, 1.5% K_4_Fe(CN)_6_ in 0.1 M cacodylate buffer. Sections were rinsed, stained with 3% uranyl acetate for 1 hour at 4°C and dehydrated in graded ethanol concentrations and propyleneoxide, followed by embedding in EMbed812. Resin blocks comprising the brain tissue were sectioned on a Leica Ultracut UCT ultramicrotome (Leica Microsystems). Semithin sections (1 mm) were collected on slides and stained with 1% Toluidine blue solution (Sigma-Aldrich) and mounted in VectaMount mounting medium (Vector). Representative areas were imaged under a Leica DM 2500M light microscope. Ultrathin sections (70 nm) were mounted on copper grids and imaged using a JEM-1400 transmission electron microscope (JEOL), equipped with an 11Mpixel Olympus SIS Quemesa camera. Images were taken at magnifications ranging from 1500X to 15000X.

### Subcellular fractionation methods

Brain mitochondria purification was adapted from the Sims method ^54^. Brains were rapidly excised, finely chopped with a scissor and homogenized with a Teflon-glass potter in 20 volumes of 20 mM Hepes, 225 mM sucrose, 75 mM mannitol, 1 mM EGTA pH 7.4, on ice. The homogenate was centrifuged at 900 g for 10 minutes at 4C to remove nuclei and unbroken debris. The supernatant (tissue homogenate) was then centrifuged at 15’000 g for 10 minutes at 4°C. The pellet is the crude brain mitochondrial fraction. The supernatant was spun at 120’000 g for 60 minutes at 4°C to remove particles and the collected supernatant was purified cytosol. The crude brain mitochondrial fraction was re-suspended in 15% Percoll and spun over a discontinuous 24/40% Percoll gradient for 6 minutes at 31,000 g at 4°C. The upper phase containing mainly myelin was discarded. A thick band in the interphase between 15/24% Percoll contains the synaptosomes. Free brain mitochondria were collected at the interphase between 24/40% Percoll. Free brain mitochondria and synaptosomes were washed in isolation buffer to remove residual Percoll.

Cultured cells were pelleted by centrifugation and re-suspended in 10 mM Hepes, sucrose 0,28 M, 1 mM EDTA buffer, pH 7.4 on ice, gently homogenized with a syringe through a 24G needle, followed by a 26G needle, and centrifuged at 800 g for 10 minutes at 4°C. The supernatant was centrifuged at 9000 g for 10 minutes at 4°C. The pellet is the enriched mitochondrial fraction.

To obtain brain nuclear enriched fraction, brain tissue was homogenized in 10 mM Hepes, 1.5 mM MgCl_2_, 10 mM KCl, 1mM dithiotreitol, pH 7.9 with a Teflon-glass tissue homogenizer on ice. Nuclei and unbroken debris were pelleted by centrifugation at 1000 g for 10 minutes at 4°C. Aliquots of 20 mM Hepes, 0.42 M NaCl, 25% (v/v) glycerol, 1.5 mM MgCl_2_, 10 mM KCl, 0.2 mM EDTA, 1mM dithiotreitol, pH 7.9 were added to the pellet and mixed by gentle mixing, followed by manual homogenization to facilitate nuclear extraction. The solution was gently mixed for 30 minutes at 4°C, and then centrifuged at 20’000 g for 10 minutes. The supernatant represents the nuclear enriched fraction.

Total tissue and cell culture lysates were obtained by lysis in radioimmunoprecipitation assay (RIPA) buffer on ice and 5 passages with a 26G syringe, followed by centrifugation at 20,000 g for 10 minutes at 4°C to remove unbroken debris.

### Immunoblot analysis

Samples were separated in reducing and denaturing conditions in NuPage gels (Invitrogen). Proteins were transferred to PVDF 0.45μm membranes, blocked with milk 5%Tris-buffered saline, Tween 20 0,1% (TTBS), and incubated with the indicated primary antibodies, washed in TTTBS incubated for 1 hour at room temperature with horseradish peroxidase conjugated secondary antibodies (Biorad) in 5% milk-TTBS, and washed in TTBS. Chemiluminescence with ECL (Amersham) and an Image Quant LAS 4000 mini was used to visualize the proteins.

### Blue native gel electrophoresis

Blue native gel electrophoresis of digitonin-solubilized mitochondria was performed as described^55^. 100 μg isolated mitochondria were solubilized with 600 μg digitonin in Invitrogen Native Page sample buffer on ice for 20 minutes, then centrifuged at 20,000 g for 20 minutes at 10 C at 4 C. 0,75% Coomassie G-250 was added to supernatants, which were loaded on a 3-12% gradient Invitrogen Native Page gel according to the instructions. After electrophoresis, mitochondrial complexes and super complexes were either visualized by protein staining with Expedeon Instant Blue or immunoblotted on a PVDF membrane and probed with the indicated antibodies.

### High-resolution respirometry

Brain mitochondria respiration was measured in Miro6 Buffer^56^ (20 mM Hepes, 110 mM sucrose, 10 mM KH_2_PO_4_, 20 mM taurine, 60 mM lactobionic acid, 3 mM MgCl2, 0.5 EGTA, pH 7.1, 1 mg/ml fatty acid free BSA, catalase 280 U/ml) at 37 °C. When needed 200 μM H_2_0_2_ was added to reoxygenate the chambers by catalase mediated O_2_ generation. 20 μg of purified free brain mitochondria or 50 μg synaptosomes were loaded into the Oroboros 2K oxygraph. A typical experiment is illustrated in Fig. 5C. Oxygen consumption rates were measured before and after addition of the following sequence of substrates and specific inhibitors: 1) 2.5 mM pyruvate, 10 mM glutamate, and 1 mM malate to measure Complex I-driven leak respiration (CI leak), 2) 2.5 mM ADP to determine complex I-driven phosphorylating respiration (CI OXPHOS). When using synaptosomes, digitonin was titrated up to 15 μg/ml to achieve synaptosomal permeabilization resulting in maximal substrate accessibility to neuronal mitochondria and maximal CI OXPHOS 3) 5 mM succinate to determine the phosphorylating respiration driven by simultaneous activation of complex I and II (CI+II OXPHOS). 4) Titrating concentrations of the mitochondrial uncoupler CCCP to reach the maximal uncoupled respiration (CI+II electron transfer capacity, ET). 5) 200 nM rotenone to fully inhibit complex I-driven respiration and measure complex II-driven uncoupled respiration (CII electron transfer capacity, CII ET). 6) 0.5 μM Antimycin A to block mitochondrial respiration at the level of complex III. Residual oxygen consumption was always negligible. 7) 2 mM ascorbate, 0.5 mM TMPD to measure cytochrome c oxidase (CIV)-driven respiration. 8) 125 μg/ml cytochrome c to evaluate mitochondrial outer membrane integrity 9) 500 μM potassium cyanide to specifically block cytochrome c oxidase activity and measure residual background oxygen consumption caused by chemical interaction between ascorbate and TMPD. Cytochrome c oxidase-driven respiration was calculated as the cyanide sensitive oxygen consumption.

### Mitochondrial respiratory chain enzyme analysis and CoQ determinations

Respiratory chain measurements were performed according to previously described protocols ^57,58^. Freeze-thawed crude brain mitochondrial fractions were resuspended in 20 mM Tris, EDTA 2 mM, pH 7.4. An aliquot was kept for CII+III activity measurements which rely on the local concentration of CoQ10 pools, while the rest was sonicated on ice with a Branson microtip sonicator at power 20%, 5 seconds ON and 20 seconds OFF for three consecutive times. Briefly, Complex I (NADH:ubiquinone oxidoreductase) activity was measured by recording the rotenone-sensitive decrease in absorbance due to oxidation of NADH at 340 nm (ε = 6.2 mM^-1^ x cm^-1^) in potassium phosphate 50 mM, pH 7.5, BSA 3 mg/ml, KCN 300 μM. The reaction was started by addition of ubiquinone. Complex II (succinate dehydrogenase) activity was measured by following the reduction of 80 μM 2,6-dichlorophenolindophenol at 600 nm (ε = 19.1 mM^-1^ x cm^-1^) in potassium phosphate 25 mM, pH 7.5, succinate 20 mM, BSA 1 mg/ml, KCN 300 μM. The reaction was started with 50μM decylubiquinone.

Complex III (ubiquinol cytochrome c oxidoreductase) activity was measured by following the Antimycin A-sensitive reduction of 75μM cytochrome c at 550 nm (ε = 18.5 mM^-1^ x cm^-1^) in potassium phosphate 25 mM, pH 7.5, tween-20 0.025%, KCN 500 μM, EDTA 100 μM. The reduction reaction was started by addition of 100μM decylubiquinol.

Complex IV (cytochrome c oxidase; EC 1.9.3.1) activity was measured using the oxidation of 60 μM reduced cytochrome c at 550 nm (ε = 18.5 mM^-1^ x cm^-1^) in potassium phosphate 50 mM, pH 7.0. The reaction was started by addition of the muscle homogenate. Complex II + III (succinate cytochrome c reductase) activity was measured using the reduction of 50 μM cytochrome c at 550 nm (ε = 18.5 mM^-1^ x cm^-1^) in 20 mM potassium phosphate, pH 7.5, succinate 10 mM, 300 μM KCN. The reaction was started by the addition of 50 μM cytochrome c. ATP synthase reverse enzymatic activity (ATP hydrolysis) was measured by recording the oligomycin-sensitive decrease in absorbance due to oxidation of NADH at 340 nm (ε = 6.2 mM^-1^ x cm^-1^) in 40 mM Tris pH 8.0, 0.2 mM EGTA, 0.2 mM NADH, 2 mM phosphoenolpyruvate, 20 mM MgCl_2_, 1 mg/ml BSA, 3 μM FCCP, 5 μg/ml Antimycin a, 4 U/ml pyruvate kinase, 4 U/ml LDH, 1 mM ATP.

Respiratory chain enzymes were normalized to the activity of the mitochondrial matrix enzyme citrate synthase. This was measured by recording the increase in absorbance at 412 nm (ε = 13.6 mM^-1^ x cm^-1^) of 5,5’-Disulfanediylbis(2-nitrobenzoic acid) DTNB 100 μM, 300μM Acetyl CoA, in Tris 100 mM, pH 8.0, 0.1% Triton X-100 after addition of 500 μM oxaloacetic acid. Citrate synthase enzymatic rates were normalized to the protein concentrations of each muscle homogenate.

CoQ content and the ratio of the reduced vs. oxidized forms were measured as described^59^.

### Antibodies

A Parl carboxy-terminal antibody was generated in house as previously reported^6^. The following commercial available antibodies were employed:

anti GFAP, Agilent (DAKO), anti IBA1 (WAKO), anti NeuN (EMD Millipore), anti-Ndufs3 (Abcam), anti-Cytocrome c (BD Phamingen), anti-Aif1 (Santa Cruz), anti-Tubulin (Abcam), anti-Caspase 3 (Cell Signaling), anti activated caspase 3 (Cell signaling), anti-Parp1 (Cell Signaling), anti-Actin (Sigma), anti-Hsp60 (BD Biosciences), ant-Uqcrfs1 (Abcam), anti-Cox4 (Abcam), anti-Sdh (Abcam), anti-Atpb (Abcam), anti-Tomm20 (Santa Cruz), anti-Vdac1 (Abcam), anti-Pink1 (Cayman), anti-Pgam5 (Sigma), anti-Ttc19 (Sigma), anti-Coq3 (Santa Cruz), anti-Coq4 (Proteintech), anti-Coq5 (Proteintech), anti-Coq6 (Proteintech), anti-Coq7 (Santa Cruz), anti-Coq9 (Santa-Cruz), anti-PDSS2 (Santa Cruz), anti-Sqrdl (Novus), anti-Ghitm (Proteintech), anti-Clpb (Proteintech), anti-Stard7 (Proteintech), anti-Smac/Diablo (Cell Signaling), anti Htra2 (R&D system), anti-Opa1 (BD Biosciences).

### Proteomic sample preparation

Mitochondria isolated from brain of *WT* and *Parl^-/-^* mice were lysed in urea buffer (8 M urea; 20 mM HEPES pH 8.0; and phosphatase inhibitors PhosStop from Roche. For each condition, triplicate samples were prepared. Samples were sonicated with a Branson 450 sonicator by 3 pulses of 10 s at an amplitude of 20% and centrifuged for 15 minutes at 16,000 g at room temperature to remove insoluble components. The protein concentration in the supernatants was measured using a BCA assay (Bio-Rad) and 500 μg total protein was used for further analysis. Proteins were reduced by adding 4.5 mM DTT and incubation for 30 minutes at 55°C. Alkylation was performed by addition of 10 mM iodoacetamide for 15 minutes in the dark at room temperature. The samples were diluted with 20 mM HEPES pH 8.0 to a urea concentration of 4 M and proteins were digested with 2 μg endoLysC (Wako) (1/250, w/w) for 4 hours at room temperature. All samples were further diluted with 20 mM HEPES pH 8.0 to a final urea concentration of 2 M and proteins were digested with 5 μg trypsin (Promega) (1/100, w/w) overnight at 37 °C. Peptides were then purified on a SampliQ SPE C18 cartridge (Agilent) and from each replicate 100 μg vacuum dried peptides were used for strong cation exchange (SCX) fractionation. To this end, SCX tips were made by stacking 3 discs (1.5 mm diameter) of polystyrene divinylbenzene copolymer with sulfonic acid (EmporeTM, 3M) in a 200 μl pipette tip. First, the SCX tips were rinsed with 100 μl acetonitrile (ACN) by pipetting the solvent in the tip, flicking the tip so the ACN touches the discs, and pushing the solvent through with a syringe. Then, each sample was dissolved in 100 μl loading buffer (1% TFA in water/ACN (95:5, v/v)) and peptides were loaded on the SCX tip. After washing the tip with 50 μl loading buffer, the peptides were fractionated by subsequently pipetting up and down 20 μl of the following fractionation buffers: 100 mM ammonium acetate, 0.5% FA (fraction 1); 175 mM ammonium acetate, 0.5% formic acid (FA) (fraction 2); 375 mM ammonium acetate, 0.5% FA (fraction 3). Remaining peptides were eluted in 2 x 20 μl of elution buffer (5% NH4OH in water/ACN (20:80, v/v)) (fraction 4). To prevent deamidation in this fraction, 2 μl 10% FA was added. Fractionated peptides were dried in a vacuum concentrator.

### LC-MS/MS and data analysis

Dried peptides from each fraction were re-dissolved in 20 μl loading solvent A (0.1% TFA in water/ACN (98:2, v/v)) of which 10 μl was injected for LC-MS/MS analysis on an Ultimate 3000 RSLCnano system in-line connected to a Q Exactive HF mass spectrometer equipped with a Nanospray Flex Ion source (Thermo). Trapping was performed at 10 μl/min for 4 min in loading solvent A on a 20 mm trapping column (made in-house, 100 μm internal diameter (I.D.), 5 μm beads, C18 Reprosil-HD, Dr. Maisch, Germany) and the sample was loaded on a 400 mm analytical column (made in-house, 75 μm I.D., 1.9 μm beads C18 Reprosil-HD, Dr. Maisch). Peptides were eluted by a non-linear increase from 2 to 56% MS solvent B (0.1% FA in water/acetonitrile (2:8, v/v)) over 140 minutes at a constant flow rate of 250 nl/min, followed by a 15-minutes wash reaching 99% MS solvent B and re-equilibration with MS solvent A (0.1% FA in water/acetonitrile (2:8, v/v)). The column temperature was kept constant at 50°C in a column oven (CoControl 3.3.05, Sonation). The mass spectrometer was operated in data-dependent mode, automatically switching between MS and MS/MS acquisition for the 16 most abundant ion peaks per MS spectrum. Full-scan MS spectra (375-1500 m/z) were acquired at a resolution of 60,000 in the orbitrap analyser after accumulation to a target value of 3,000,000. The 16 most intense ions above a threshold value of 13,000 were isolated (window of 1.5 Th) for fragmentation at a normalized collision energy of 32% after filling the trap at a target value of 100,000 for maximum 80 ms. MS/MS spectra (200-2000 m/z) were acquired at a resolution of 15,000 in the orbitrap analyser. The S-lens RF level was set at 55 and we excluded precursor ions with single and unassigned charge states from fragmentation selection.

Data analysis was performed with MaxQuant (version 1.5.3.30) using the Andromeda search engine with default search settings including a false discovery rate set at 1% on both the peptide and protein level. Spectra were searched against the mouse proteins in the UniProt/Swiss-Prot database (database release version of March 2016 containing 16,618 mouse protein sequences) with the mass tolerance for precursor and fragment ions set to 4.5 and 20 ppm, respectively, during the main search. Enzyme specificity was set as C-terminal to arginine and lysine (trypsin), also allowing cleavage at arginine/lysine-proline bonds with a maximum of two missed cleavages. Carbamidomethylation of cysteine residues was set as a fixed modification and variable modifications were set to oxidation of methionine residues (to sulfoxides) and acetylation of protein N-termini. Proteins were quantified by the MaxLFQ algorithm integrated in the MaxQuant software. Only proteins with at least one unique or razor peptide were retained for identification, while a minimum ratio count of two unique or razor peptides was required for quantification. Further data analysis was performed with the Perseus software (version 1.5.2.6) after loading the protein groups file from MaxQuant. Proteins were only identified by site, reverse database hits, and potential contaminants were removed and replicate samples were grouped. Proteins with less than three valid values in at least one group were removed and missing values were imputed from a normal distribution around the detection limit. In this way, a total of 2,154 proteins were quantified. Then, a t-test was performed (FDR=0.05 and S0=0.5) to reveal proteins that are significantly different between WT and *Parl*^-/-^ samples and to generate the volcano plot depicted in Fig.6A.

### Plasmids

All mouse Parl mutants were constructed using the QuickChange II XL mutagenesis kit (Stratagene). Immortalized mouse embryonic fibroblasts (MEFs) derived from *Parl*^-/-^ mice were cultured in Dulbecco’s modified Eagle’s medium/F-12 (Gibco) containing 10% fetal bovine serum (Gibco). At 30–40% confluence, the MEFs were transduced using a replication-defective recombinant retroviral expression system (Clontech) with either wild-type (Parl^WT^) or with catalytic inactive Parl^S275A^. Cell lines stably expressing the desired proteins were selected based on their acquired resistance to 5 μg/ml puromycin.

### mtDNA quantification

Isolation of DNA was done using the Qiagen Dneasy Blood and Tissue kit following the manufacturers instructions. Real-time semi-quantitative PCR was performed on a LightCycler 480 (Roche) using the SensiFast Sybr No ROX Mix (Bioline) according to the manufacturers instructions. In each reaction 1,5ng of gDNA was used. Sequences of the primers used can be found in the table below. Crossing points were determined by using the second derivative method. Fold changes were calculated with the ΔΔCt method^60^

~~~
COX1 forward      TCGCCATCATATTCGTAGGAG
COX1 reverse      GTAGCGTCGTGGTATTCCTGA
ND4 forward       TTATTACCCGATGAGGGAACC
ND4 reverse       GAGGGCAATTAGCAGTGGAAT
ApoB forward      CGTGGGCTCCAGCATTCTA
ApoB reverse     TCACCAGTCATTTCTGCCTTT
B2m forward      TGTCAGATATGTCCTTCAGCAAGG
B2m reverse     TGCTTAACTCTGCAGGCGTATG
~~~

### Cell culture

Mouse primary neurons were cultured using a protocol adapted from Goslin and Banker^61^. Briefly, E14,5 mouse whole brains were trypsinized, dissociated with fire-polished Pasteur pipets, and plated onto poly-L-lysine (Sigma)-coated culture dishes. Cells were incubated for the first 4 hr in minimum essential medium (MEM) with 10% horse serum and then maintained in serum-free neurobasal medium with B7 supplement (Gibco). All cells were incubated at 37°C with 5% CO2. When specified, after one week in culture, neurons were treated with 10μM etoposide for 24h before collection.

### Statistical analysis

All numerical data are expressed as mean ± SD. Two sided student’s t test was used to compare differences between two groups. Statistical significance for survival curves was calculated by log-rank (Mantel-Cox). Differences were considered statistical significant for p ≤ 0.05. Mass spectrometry data were analyzed as detailed in the LC-MS/MS and data analysis section.

## ACKNOWLEDGEMENTS

This work was supported by the Fonds voor Wetenschappelijk Onderzoek (FWO), the KU Leuven and VIB, a Methusalem grant from the KU Leuven/Flemish Government to BDS. BDS is supported by the Bax-Vanluffelen Chair for Alzheimer’s Disease and “Opening the Future” of the Leuven Universiteit Fonds (LUF). This work is supported by Vlaams Initiatief voor Netwerken voor Dementie Onderzoek (VIND, Strategic Basic Research Grant 135043). This work was partially supported by Spanish Ministry of Health, Instituto de Salud Carlos III (ISCIII), FIS PI17-01286 grant. Confocal microscope equipment was acquired through a Hercules Type 1 AKUL/09/037 to W. Annaert. Confocal and Electron microscopy were performed in the VIB Bio Imaging Core (LiMoNe and EMoNe facilities). We are grateful to A. Francis and E. Seuntjens for ISH through InfraMouse (Hercules Foundation type 3 large infrastructure ZW09-03). We thank Veronique Hendrickx and Jonas Verwaeren for help with the mouse colonies. M.S. is the recipient of an EMBO long-term fellowship (ALTF 648-2013).

## REFERENCES

1. Pellegrini, L. et al. PAMP and PARL, two novel putative metalloproteases interacting with the COOH-terminus of Presenilin-1 and -2. J. Alzheimers. Dis. 3, 181–190 (2001).

2. Spinazzi, M. & De Strooper, B. PARL: The mitochondrial rhomboid protease. Semin. Cell Dev. Biol. 60, 19–28 (2016).

3. Düsterhöft, S., Künzel, U. & Freeman, M. Rhomboid proteases in human disease: Mechanisms and future prospects. Biochim. Biophys. Acta - Mol. Cell Res. 0–1 (2017). doi:10.1016/j.bbamcr.2017.04.016

4. Urban, S. A guide to the rhomboid protein superfamily in development and disease. Semin. Cell Dev. Biol. 60, 1–4 (2016).

5. Shi, G. et al. Functional alteration of PARL contributes to mitochondrial dysregulation in Parkinson’s disease. Hum. Mol. Genet. 20, 1966–1974 (2011).

6. Cipolat, S. et al. Mitochondrial Rhomboid PARL Regulates Cytochrome c Release during Apoptosis via OPA1-Dependent Cristae Remodeling. Cell 126, 163–175 (2006).

7. Anand, R. et al. The i-AAA protease YME1L and OMA1 cleave OPA1 to balance mitochondrial fusion and fission. J. Cell Biol. 204, 919–929 (2014).

8. Griparic, L., Kanazawa, T. & Van Der Bliek, A. M. Regulation of the mitochondrial dynamin-like protein Opa1 by proteolytic cleavage. J. Cell Biol. 178, 757–764 (2007).

9. Chao, J.-R. et al. Hax1-mediated processing of HtrA2 by Parl allows survival of lymphocytes and neurons. Nature 452, 98–102 (2008).

10. Deas, E. et al. PINK1 cleavage at position A103 by the mitochondrial protease PARL. Hum. Mol. Genet. 20, 867–879 (2011).

11. Jin, S. M. et al. Mitochondrial membrane potential regulates PINK1 import and proteolytic destabilization by PARL. J. Cell Biol. 191, 933–942 (2010).

12. Meissner, C., Lorenz, H., Weihofen, A., Selkoe, D. J. & Lemberg, M. K. The mitochondrial intramembrane protease PARL cleaves human Pink1 to regulate Pink1 trafficking. J. Neurochem. 117, 856–867 (2011).

13. Sekine, S. et al. Rhomboid protease PARL mediates the mitochondrial membrane potential loss-induced cleavage of PGAM5. J. Biol. Chem. 287, 34635–34645 (2012).

14. Saita, S. et al. PARL mediates Smac proteolytic maturation in mitochondria to promote apoptosis. Nat. Cell Biol. 19, 318–328 (2017).

15. Valente, E. M. et al. Hereditary Early-Onset Parkinson ’ s Disease Caused by Mutations in PINK1. Science (80-. ). 304, 1158–1161 (2004).

16. Lu, W. et al. Genetic deficiency of the mitochondrial protein PGAM5 causes a Parkinson’s-like movement disorder. Nat. Commun. 5, 4930 (2014).

17. Greene, A. W. et al. Mitochondrial processing peptidase regulates PINK1 processing, import and Parkin recruitment. EMBO Rep. 13, 378–385 (2012).

18. Civitarese, A. E. et al. Regulation of skeletal muscle oxidative capacity and insulin signaling by the mitochondrial rhomboid protease PARL. Cell metab 11, 412–426 (2010).

19. Jeyaraju, D. V et al. Phosphorylation and cleavage of presenilin-associated rhomboid-like protein (PARL) promotes changes in mitochondrial morphology. Proc. Natl. Acad. Sci. U. S. A. 103, 18562–18567 (2006).

20. Meissner, C., Lorenz, H., Hehn, B. & Lemberg, M. K. Intramembrane protease PARL defines a negative regulator of PINK1- and PARK2/Parkin-dependent mitophagy. Autophagy 11, 1484–1498 (2015).

21. Shi, G. & McQuibban, G. A. The Mitochondrial Rhomboid Protease PARL Is Regulated by PDK2 to Integrate Mitochondrial Quality Control and Metabolism. Cell Rep. 18, 1458–1472 (2017).

22. Lake, N. J., Compton, A. G., Rahman, S. & Thorburn, D. R. Leigh syndrome: One disorder, more than 75 monogenic causes. Ann. Neurol. 79, 190–203 (2016).

23. Lake, N. J., Bird, M. J., Isohanni, P. & Paetau, A. Leigh syndrome: neuropathology and pathogenesis. J. Neuropathol. Exp. Neurol. 74, 482–92 (2015).

24. Saita, S., Tatsuta, T., Lampe, P. A. & König, T. PARL partitions the lipid transfer protein STARD7 between the cytosol and mitochondria. 1–18 (2018). doi:10.15252/embj.201797909

25. Youle, R. J. & Narendra, D. P. Mechanisms of mitophagy. Nat Rev Mol Cell Biol 12, 9–14 (2011).

26. Wang, Z., Jiang, H., Chen, S., Du, F. & Wang, X. The mitochondrial phosphatase PGAM5 functions at the convergence point of multiple necrotic death pathways. Cell 148, 228–243 (2012).

27. Fernández-Vizarra, E., Tiranti, V. & Zeviani, M. Assembly of the oxidative phosphorylation system in humans: What we have learned by studying its defects. Biochim. Biophys. Acta - Mol. Cell Res. 1793, 200–211 (2009).

28. Bottani, E. et al. TTC19 Plays a Husbandry Role on UQCRFS1 Turnover in the Biogenesis of Mitochondrial Respiratory Complex III Article TTC19 Plays a Husbandry Role on UQCRFS1 Turnover in the Biogenesis of Mitochondrial Respiratory Complex III. Mol. Cell 1–10 (2017). doi:10.1016/j.molcel.2017.06.001

29. Koch, J. et al. Mutations in TTC19: Expanding the molecular, clinical and biochemical phenotype. Orphanet J. Rare Dis. 10, 1–12 (2015).

30. Guarás, A. et al. The CoQH2/CoQ Ratio Serves as a Sensor of Respiratory Chain Efficiency. Cell Rep. 15, 197–209 (2016).

31. Ghezzi, D. et al. Mutations in TTC19 cause mitochondrial complex III deficiency and neurological impairment in humans and flies. Nat. Genet. 43, 259–63 (2011).

32. Ziosi, M. et al. Coenzyme Q deficiency causes impairment of the sulfide oxidation pathway. EMBO Mol. Med. 9, 96–111 (2017).

33. Kruse, S. E. et al. Mice with Mitochondrial Complex I Deficiency Develop a Fatal Encephalomyopathy. Cell Metab. 7, 312–320 (2008).

34. Quintana, A., Kruse, S. E., Kapur, R. P., Sanz, E. & Palmiter, R. D. Complex I deficiency due to loss of Ndufs4 in the brain results in progressive encephalopathy resembling Leigh syndrome. Proc. Natl. Acad. Sci. U. S. A. 107, 10996–1001 (2010).

35. McQuibban, G. A., Lee, J. R., Zheng, L., Juusola, M. & Freeman, M. Normal Mitochondrial Dynamics Requires Rhomboid-7 and Affects Drosophila Lifespan and Neuronal Function. Curr. Biol. 16, 982–989 (2006).

36. Patterson, V. L. et al. Neural-specific deletion of HTRA2 causes cerebellar neurodegeneration and defective processing of mitochondrial OPA1. PLoS One 9, 1–24 (2014).

37. Dooley, J. & Liston, A. Molecular control over thymic involution: From cytokines and microRNA to aging and adipose tissue. Eur. J. Immunol. 42, 1073–1079 (2012).

38. Savino, W. The thymus gland is a target in malnutrition. Eur. J. Clin. Nutr. 56, S46–S49 (2002).

39. Seksenyan, A. et al. Thymic involution, a co-morbidity factor in amyotrophic lateral sclerosis. J. Cell. Mol. Med. 14, 2470–2482 (2010).

40. Offner, H. et al. Increased Regulatory T Cells and Circulating Macrophages 1. J. Immunol. 176, 6523–6531 (2016).

41. Jeyaraju, D. V, Cisbani, G., De Brito, O. M., Koonin, E. V & Pellegrini, L. Hax1 lacks BH modules and is peripherally associated to heavy membranes: implications for Omi/HtrA2 and PARL activity in the regulation of mitochondrial stress and apoptosis. Cell Death Differ. 16, 1622–1629 (2009).

42. Acosta, M. J. et al. Coenzyme Q biosynthesis in health and disease. Biochim. Biophys. Acta - Bioenerg. 1857, 1079–1085 (2016).

43. Luna-Sánchez, M. et al. CoQ deficiency causes disruption of mitochondrial sulfide oxidation, a new pathomechanism associated with this syndrome. EMBO Mol. Med. 9, 78–95 (2017).

44. Alcázar-Fabra, M., Navas, P. & Brea-Calvo, G. Coenzyme Q biosynthesis and its role in the respiratory chain structure. Biochim. Biophys. Acta - Bioenerg. 1857, 1073–1078 (2016).

45. Stefely, J. A. & Pagliarini, D. J. Biochemistry of Mitochondrial Coenzyme Q Biosynthesis. Trends Biochem. Sci. 42, 824–843 (2017).

46. Veling, M. T. et al. Multi-omic Mitoprotease Profiling Defines a Role for Oct1p in Coenzyme Q Production. Mol. Cell 68, 970–977.e11 (2017).

47. Kühl, I. et al. Transcriptomic and proteomic landscape of mitochondrial dysfunction reveals secondary coenzyme Q deficiency in mammals. Elife 6, e30952 (2017).

48. Salviati, L. et al. Haploinsufficiency of COQ4 causes coenzyme Q10deficiency. J. Med. Genet. 49, 187–191 (2012).

49. Tronche, F. et al. Disruption of the glucocorticoid receptor gene in the nervous system results in reduced anxiety. Nat. Genet. 23, 99–103 (1999).

50. Brüning, J. C. et al. A muscle-specific insulin receptor knockout exhibits features of the metabolic syndrome of NIDDM without altering glucose tolerance. Mol. Cell 2, 559–69 (1998).

51. Morais, V. A. et al. Parkinson’s disease mutations in PINK1 result in decreased Complex I activity and deficient synaptic function. EMBO Mol. Med. 1, 99–111 (2009).

52. Kaufmann, W. et al. Proliferative and Nonproliferative Lesions of the Rat and Mouse Central and Peripheral Nervous Systems. Toxicol. Pathol. 40, 87S–157S (2012).

53. Seuntjens, E. et al. Sip1 regulates sequential fate decisions by feedback signaling from postmitotic neurons to progenitors. Nat. Neurosci. 12, 1373–1380 (2009).

54. Sims, N. R. & Anderson, M. F. Isolation of mitochondria from rat brain using Percoll density gradient centrifugation. Nat. Protoc. 3, 1228–1239 (2008).

55. Jha, P., Wang, X. & Auwerx, J. Analysis of Mitochondrial Respiratory Chain Supercomplexes Using Blue Native Polyacrylamide Gel Electrophoresis (BN-PAGE). Curr. Protoc. Mouse Biol. 6, 1–14 (2016).

56. Fasching, M., Renner-sattler, K. & Gnaiger, E. Mitochondrial Respiration Medium - MiR06. Mitochondrial Physiol. Netw. 14.13, 1–4 (2016).

57. Spinazzi, M., Casarin, A., Pertegato, V., Salviati, L. & Angelini, C. Assessment of mitochondrial respiratory chain enzymatic activities on tissues and cultured cells. Nat. Protoc. 7, 1235–1246 (2012).

58. Spinazzi, M. et al. Optimization of respiratory chain enzymatic assays in muscle for the diagnosis of mitochondrial disorders. Mitochondrion 11, 893–904 (2011).

59. Rodríguez-Aguilera, J., Cortés, A., Fernández-Ayala, D. & Navas, P. Biochemical Assessment of Coenzyme Q10 Deficiency. J. Clin. Med. 6, 27 (2017).

60. Livak, K. J. & Schmittgen, T. D. Analysis of relative gene expression data using real-time quantitative PCR and the 2-ΔΔCT method. Methods 25, 402–408 (2001).

61. Goslin, K. & Banker, G. Rapid changes in the distribution of GAP-43 correlate with the expression of neuronal polarity during normal development and under experimental conditions. J.Cell Biol. 110, 1319–1331 (1990).

